# The lysosomal chloride-proton exchanger CLC7 functions in melanosomes as a negative regulator of human pigmentation

**DOI:** 10.1101/2021.02.05.430016

**Authors:** Donald C. Koroma, Jessica L. Scales, Joshaya C. Trotman, Kazumasa Wakamatsu, Shosuke Ito, Elena Oancea

**Author notes:** Address correspondence to: Elena Oancea.

## Abstract

Mutations in the Cl^−^/H^+^ exchanger CLC7 and its subunit OSTM1 result in osteopetrosis, lysosomal disorders, and pigmentation defects in mice and humans. How CLC7/OSTM1 regulates pigmentation in skin and hair melanocytes remains unexplored. In human epidermal melanocytes, we found CLC7/OSTM1 localized to melanosomes, the organelles in which melanin is synthesized, where it negatively regulates melanin production. Using a novel ratiometric melanosomal pH indicator, we showed that CLC7 acidifies melanosomes, opposing the function of the oculocutaneous albinism II (OCA2) Cl^−^ ion channel. The *de novo* CLC7 variant (CLC7-Y715C) that causes albinism in humans and mice, decreased melanocytes pigmentation, which was restored by coexpression of OCA2. Remarkably, the enlarged hyperacidic vacuoles caused by CLC7-Y715C were also rescued by OCA2 coexpression in both melanocytes and non-melanocytic cells. Our data uncover a novel mechanism by which CLC7 regulates melanocyte pigmentation and identifies OCA2 as a tool to counteract the effects of CLC7 activating mutations.

## Introduction

Melanosomes are specialized lysosome-related organelles in melanocytes that synthesize and store melanin pigments responsible for skin, hair, and eye pigmentation (1, 2). The synthesis of melanin involves the concerted action of melanogenic enzymes that function optimally in a luminal environment with a near neutral pH (3, 4). Maintenance of melanosomal pH requires the movement of ions across the melanosome membrane via resident ion channels and transporters, and is crucial for melanin synthesis (5–7). Indeed, mutations in several genes encoding ion transport proteins in melanosomes are associated with pigmentation disorders like oculocutaneous albinism (OCA) (8–10). Despite the importance of melanosome function in pigmentation and vision, the proteins and mechanisms that regulate ionic and pH homeostasis in melanosomes remain poorly understood.

Recently, we used a comparative analysis of the human epidermal melanocyte and keratinocyte transcriptome (11, 12) to identify melanocyte-enriched genes encoding known or putative melanosomal ion channels and transporters (13). Among these, the chloride/proton (Cl^−^/H^+^) exchanger Chloride Channel 7 (CLC7) and its obligate β-subunit Osteopetrosis-Associated Transmembrane Protein 1 (OSTM1) were highly enriched in epidermal melanocytes compared to keratinocytes (13). Interestingly, CLC7^−/−^ mice have grey fur in an agouti background (14, 15), a coat color phenotype also shared by the OSTM1^−/−^ *grey-lethal* mice (16, 17), demonstrating that both CLC7 and OSTM1 regulate mouse fur pigmentation, but via unknown mechanisms. More recently, a pathogenic *de novo* point mutation in the CLC7 gene (Y715C) was identified in two patients with lysosomal storage defects, developmental delays, and hypopigmentation (18), suggesting that CLC7 also regulates human skin pigmentation, albeit through a mechanism that remains elusive.

CLC7 and OSTM1 function in lysosomes as a 2Cl^−^/1H^+^ exchanger and provide the main lysosomal permeation pathway for Cl^−^ (19). CLC7 also contributes to lysosomal acidification (19, 20), although this function remains a matter of debate (15, 21), partly due to potential compensatory mechanisms and a lack of reliable and accurate ratiometric pH sensors (15, 20, 22). In lysosomes, CLC7 provides a parallel shunt conductance for the electrogenic vacuolar H^+^-ATPase (V-ATPase), which requires an influx of counterions like Cl^−^ to facilitate net H^+^ influx and luminal acidification (20, 22–24). Interestingly, the pathogenic *de novo* CLC7-Y715C mutation enhances CLC7-mediated Cl^−^ flux (gain-of-function), resulting in hyperacidic lysosomes and accumulation of large intracellular vacuoles, both consistent with defective lysosomal storage (18).

Melanosomes are derived from organelles of the endocytic lineage, but mature as specialized organelles that synthesize melanin in a process that is highly reliant on luminal pH. In this study, we sought to identify the mechanism by which CLC7 regulates melanocyte pigmentation. We found that in human epidermal melanocytes (HEMs), CLC7 localizes to both melanosomes and lysosomes, and regulates not only the amount, but also the type of melanin pigment synthesized. We designed a novel genetically-encoded and ratiometric pH indicator for melanosomes, RpHIMEL, and employed it to show that CLC7 acidifies melanosomes, thus inhibiting melanin synthesis in human pigmented cells. Interestingly, CLC7 localizes to the same population of melanosomes as the Cl^−^ ion channel OCA2 (6) and opposes its function: ectopic expression of OCA2 rescues the phenotype of the melanocytes expressing the CLC7-Y715C gain-of-function mutant. Remarkably, in non-melanocytic cells, OCA2 localizes to lysosomes and rescues the phenotype of CLC7-Y715C expressing cells, suggesting that OCA2 might represent a therapeutic target for CLC7-Y715C patient cells.

## Results

### CLC7 and OSTM1 are negative regulators of pigmentation in melanocytes

We identified CLC7 and its obligate β-subunit OSTM1 based on transcriptomic analyses for transmembrane proteins that are differentially expressed in human epidermal melanocytes (HEMs) compared to keratinocytes (13). To test if these candidate genes regulate pigmentation, we optimized a short hairpin RNA (shRNA) screen to examine the effect of CLC7 and OSTM1 downregulation on cellular melanin concentration in MNT1 pigmented human melanoma cells (13), an established cell model for pigmentation studies (25, 26). Expression of two distinct CLC7-targeting shRNAs (#1 or #2) led to increased pigmentation of MNT1 cells compared to control cells expressing non-targeting (NT) shRNA (**Fig. S1A**). MNT1 cells expressing CLC7 shRNA #1 or shRNA #2 had 2.5-fold and 8.0-fold higher cellular melanin, respectively, compared to control NT shRNA cells (**Fig. S1B**), in agreement with lower CLC7 mRNA and protein expression (**Fig. S1C, S1D**). As the downregulation of CLC7 expression was more robust in cells expressing CLC7 shRNA #2, we used this shRNA for further experiments.

We next tested if reducing the expression of CLC7 had a similar effect on pigmentation in primary human melanocytes (HEMs). We measured the effect of CLC7 shRNA on CLC7 expression in HEMs and found that CLC7 mRNA and protein levels were reduced by 84% and 94%, respectively (**Fig. 1B, 1C**, green bars). CLC7 shRNA also reduced OSTM1 mRNA levels by 74% compared to control NT shRNA cells (**Fig. 1B**, blue bars). As expected, OSTM1 shRNA reduced OSTM1 mRNA levels by 81% (**Fig. 1B**, blue bars). Although OSTM1 shRNA did not significantly alter CLC7 mRNA expression (**Fig. 1B**), it decreased CLC7 protein levels by 59% (**Fig. 1C, S1E**), consistent with previous observations in OSTM1 and CLC7 mutant mice (27). HEMs expressing shRNA targeting CLC7 or its β-subunit OSTM1 had visibly darker pigmentation and more than a 2-fold increase in cellular melanin concentration (**Fig. 1A, 1D**). We wondered if the observed effect on pigmentation could be due to upregulation of the other two intracellular CLC isoforms: CLC3 and CLC5, both expressed in HEMs, albeit at significantly lower levels than CLC7 (Fig. **S2A**), consistent with RNA-Seq data (11, 12). CLC3 and CLC5 mRNA and protein levels were not significantly altered in HEMs expressing CLC7-targeting shRNA (Fig. **S2B-E**), suggesting that the changes in melanocyte pigmentation are caused by decreased CLC7 expression.

**Figure 1.**
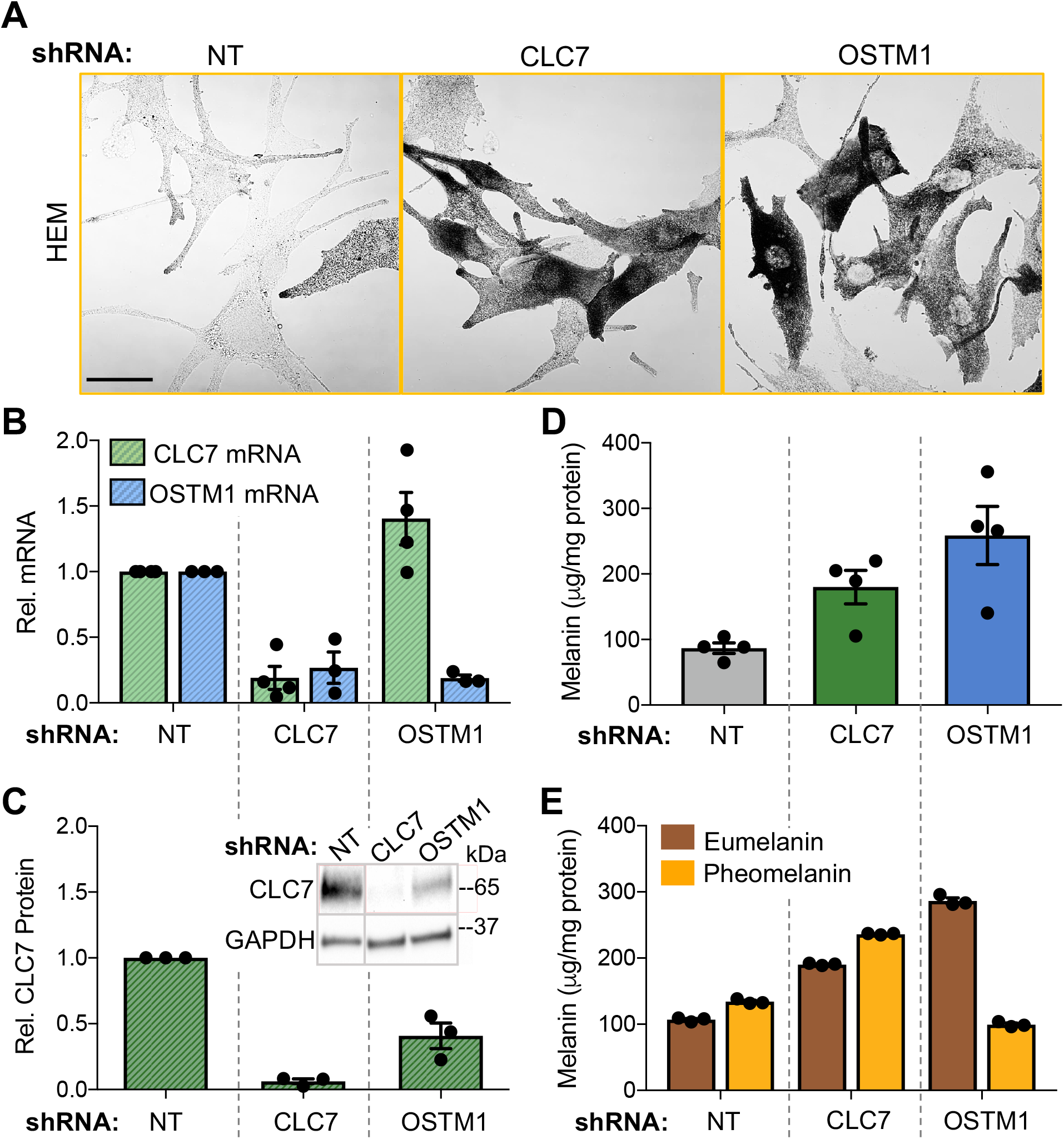
Downregulation of CLC7 increases pigmentation in primary human melanocytes. **A.** Representative bright field images of HEMs expressing CLC7− and OSTM1-targeting shRNA show increased pigmentation compared to HEMs expressing control non-targeting (NT) shRNA. Scale bar: 20 μm. **B.** CLC7 mRNA levels (green bars) were reduced by 84% in HEMs expressing CLC7 shRNA (P = 0.0027) and slightly increased in cells expressing OSTM1 shRNA (P = 0.1367) compared to control NT shRNA cells. OSTM1 mRNA levels (blue bars) were reduced by 81% in OSTM1 shRNA expressing HEMs (P = 0.0009) and by 74% in HEMs expressing CLC7 shRNA (P = 0.0256) compared to NT shRNA cells. P-values derived from paired two-sample t-tests. ± SEM; n ≥ 3 independent experiments per condition. **C.** Compared to HEMs expressing NT shRNA, CLC7 protein levels were decreased by 94% in CLC7 shRNA cells (P = 0.0004) and by 59% in OSTM1 shRNA cells (P = 0.0257) *Inset:* Representative immunoblot showing attenuated CLC7 protein levels in HEMs expressing CLC7 or OSTM1 shRNA. Glyceraldehyde 3-phosphate dehydrogenase (GAPDH) served as a loading control. P-values derived from paired two-sample t-tests. ± SEM; n = 3 independent experiments. **D.** Total cellular melanin concentration is increased 2.1-fold in HEMs expressing CLC7-targeting shRNA (green bar; P = 0.0161) and 3.0-fold in OSTM1-targeting shRNA expressing HEMs (blue bar; P = 0.0178), compared to NT shRNA cells. P-values derived from paired two-sample t-tests. ± SEM; n = 4 independent experiments. **E.** Total cellular concentration of eumelanin (measured as PTCA, see Methods; brown bars) is increased in HEMs expressing CLC7- or OSTM1-targeting compared to NT shRNA (P = 0.0001 and 0.0004, respectively). The total cellular concentration of pheomelanin (measured as BZ-PM + BT-PM, see Methods; orange bars) is increased by 76% in CLC7 shRNA cells (P = 0.0004) and decreased by 26% in OSTM1 shRNA cells (P = 0.0006). P-values derived from paired two-sample t-tests. ± SEM; n = 3 independent experiments.

Two types of melanin pigments are synthesized in human skin and mouse hair: the yellow-red pheomelanin and the brown-black eumelanin (28). Our results showing increased pigmentation in melanocytes with reduced CLC7 or OSTM1 expression (**Fig. 1C**) are in contrast to the coat-color phenotype of CLC7^−/−^ and OSTM1^−/−^ mice, both of which have a grey coat color due to reduced pheomelanin (15, 16). To determine how downregulation of CLC7 or OSTM1 affects each type of melanin in human skin melanocytes, we analyzed by high-performance liquid chromatography (HPLC) extracts from HEMs stably expressing CLC7- or OSTM1-targeting shRNA (28-30) and found that, unlike in mouse hair, both eumelanin and pheomelanin concentration increased with CLC7 downregulation (**Fig. 1E**). In contrast, reduced expression of OSTM1 led to higher eumelanin, but lower pheomelanin concentrations in HEMs (**Fig. 1E**), similar to the mouse grey coat phenotype. These measurements account for the increase in total melanin measured in **Fig. 1D** and suggest that CLC7/OSTM1 regulate pigmentation by different mechanisms in human skin and mouse hair.

### CLC7 localizes to lysosomes and melanosomes in pigmented cells

In non-melanocytic cells, CLC7 and OSTM1 function as a Cl^−^/H^+^ antiporter at lysosomal membranes (19, 31, 32). We sought to identify CLC7 and OSTM1 localization in pigmented cells. Co-immunostaining of HEMs for endogenous CLC7 and lysosomal-associated membrane protein 1 (LAMP1) resulted in 49% overlap, suggesting that approximately half of CLC7 protein localizes to lysosomes (**Fig. 2A, 2B**, top). Interestingly, CLC7 and the melanosomal marker tyrosinase-related protein 1 (TYRP1) had a 45% overlap, suggesting that approximately half of CLC7 localizes to melanosomes (**Fig. 2A, 2B**, bottom). Although LAMP1 and TYRP1 could be found together in a small population of melanosomes (<10%, (33)), the majority of LAMP1 and TYRP1 represent different populations of organelles. Our data suggest that in melanocytes, CLC7’s functions may be equally divided between lysosomes and melanosomes. To further investigate CLC7 localization and function in melanosomes, we ectopically expressed in HEMs fluorescently-tagged and functional GFP-CLC7 together with OSTM1-Myc, which overlapped with endogenous CLC7 (**Fig. S3A, S3B**). Approximately half of GFP-CLC7 colocalized with either LAMP1- and TYRP1-positive organelles, similar to endogenous CLC7 (**Fig. 2C, 2D**).

**Figure 2.**
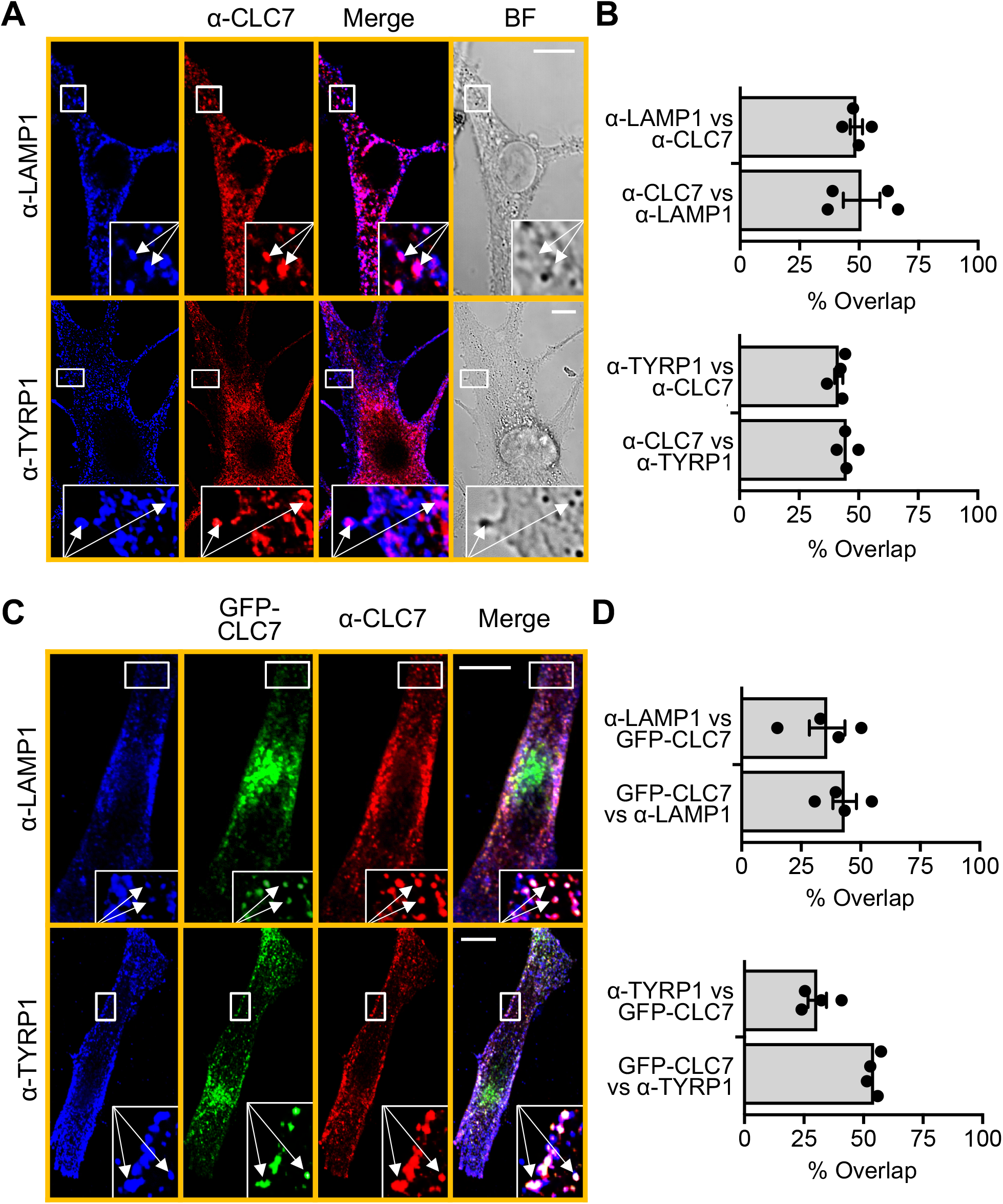
CLC7 localizes to both melanosomes and lysosomes. **A.** Representative confocal images of HEMs immunostained with antibodies to lysosome-associated membrane protein 1 (LAMP1; top) and tyrosinase related protein 1 (TYRP1; bottom) to visualize lysosomes and melanosomes, respectively. Immunostaining reveals CLC7 colocalization with LAMP1- and TYRP1-stained organelles and darkly pigmented melanosomes observed by bright field (BF) microscopy. *Inset:* Magnified view of areas in the white boxes. Arrows indicate organelles with endogenous CLC7. Scale bar: 10 μm. **B.** Quantification of images as in **A**, showing reciprocal overlap between endogenous CLC7 and LAMP1 (top) or TYRP1 (bottom) in HEMs. Endogenous CLC7 localizes to 49% and 45% of all LAMP1- and TYRP1-stained organelles, respectively. LAMP1- and TYRP1-stained organelles colocalize with 51% and 42% of total endogenous CLC7, respectively. ± SEM; n = 4 independent experiments. **C.** Representative confocal images of HEMs transiently transfected with GFP-CLC7 and OSTM1-Myc and immunostained with antibodies to LAMP1 (top) and TYRP1 (bottom). Similar to endogenous CLC7, overexpressed GFP-CLC7 localizes to LAMP1- and TYRP1-stained organelles. *Inset:* Magnified view of areas in the white boxes. Arrows indicate stained organelles with GFP-CLC7. Scale bar: 10 μm. **D.** Quantification of images as in **C** showing reciprocal overlap between overexpressed GFP-CLC7 and LAMP1 (top) or TYRP1 (bottom) in HEMs. GFP-CLC7 localizes to 36% and 31% of all LAMP1- and TYRP1-stained organelles, respectively. LAMP1- and TYRP1-stained organelles colocalize with 43% and 54% of total GFP-CLC7, respectively. ± SEM; n = 4 independent experiments.

### CLC7 colocalizes with the melanosomal Cl^−^-selective ion channel OCA2

We next investigated the mechanism by which melanosomal CLC7 might regulate pigmentation. In lysosomes, CLC7 facilitates the inward transport of 2Cl^−^ in exchange for outward transport of each H^+^ (19). OCA2 positively regulates pigmentation via the outward transport of Cl^−^ from the melanosome lumen, which decreases the driving force for inward H^+^ transport by V-ATPase, leading to a more neutral pH, favorable for melanin synthesis (**Fig. 3A**). However, OCA2-mediated Cl^−^ flux is dependent on the Cl^−^ concentration ([Cl^−^]) gradient across the melanosome membrane ([Cl^−^]_lumen_ > [Cl^−^]_cytosol_), suggesting that OCA2 function requires maintenance of the [Cl^−^] gradient by Cl^−^-selective melanosomal transporters. To identify if CLC7 could serve such function, we first determined whether CLC7 and OCA2 are found in the same population of melanosomes. In the absence of specific OCA2 antibodies, we ectopically expressed GFP-CLC7 and OSTM1-Myc together with mCherry (mCh)-OCA2 in HEMs and MNT1 cells, and found greater than 60% overlap between GFP-CLC7 and mCh-OCA2 in both cell types (**Fig. 3B, 3C**), suggesting that both CLC7 and OCA2 are present in a populations of early-stage melanosomes (34). Because CLC7 and OCA2 have opposing effects on pigmentation, we sought to determine if melanin synthesis is dually regulated by CLC7 and OCA2. In MNT1 cells, ectopic expression of mCh-OCA2 increased cellular melanin by 1.5-fold (**Fig. 3D**), while GFP-CLC7 expression decreased cellular melanin by 1.8-fold, consistent with pigmentation being positively regulated by OCA2 (6) and negatively regulated by CLC7 (**Fig. 1**). Interestingly, coexpression of OCA2 with CLC7 and OSTM1 restored cellular melanin to levels observed in control MNT1 cells (**Fig. 3D**), suggesting that both CLC7 and OCA2 might regulate melanin synthesis by modulating melanosomal pH.

**Figure 3.**
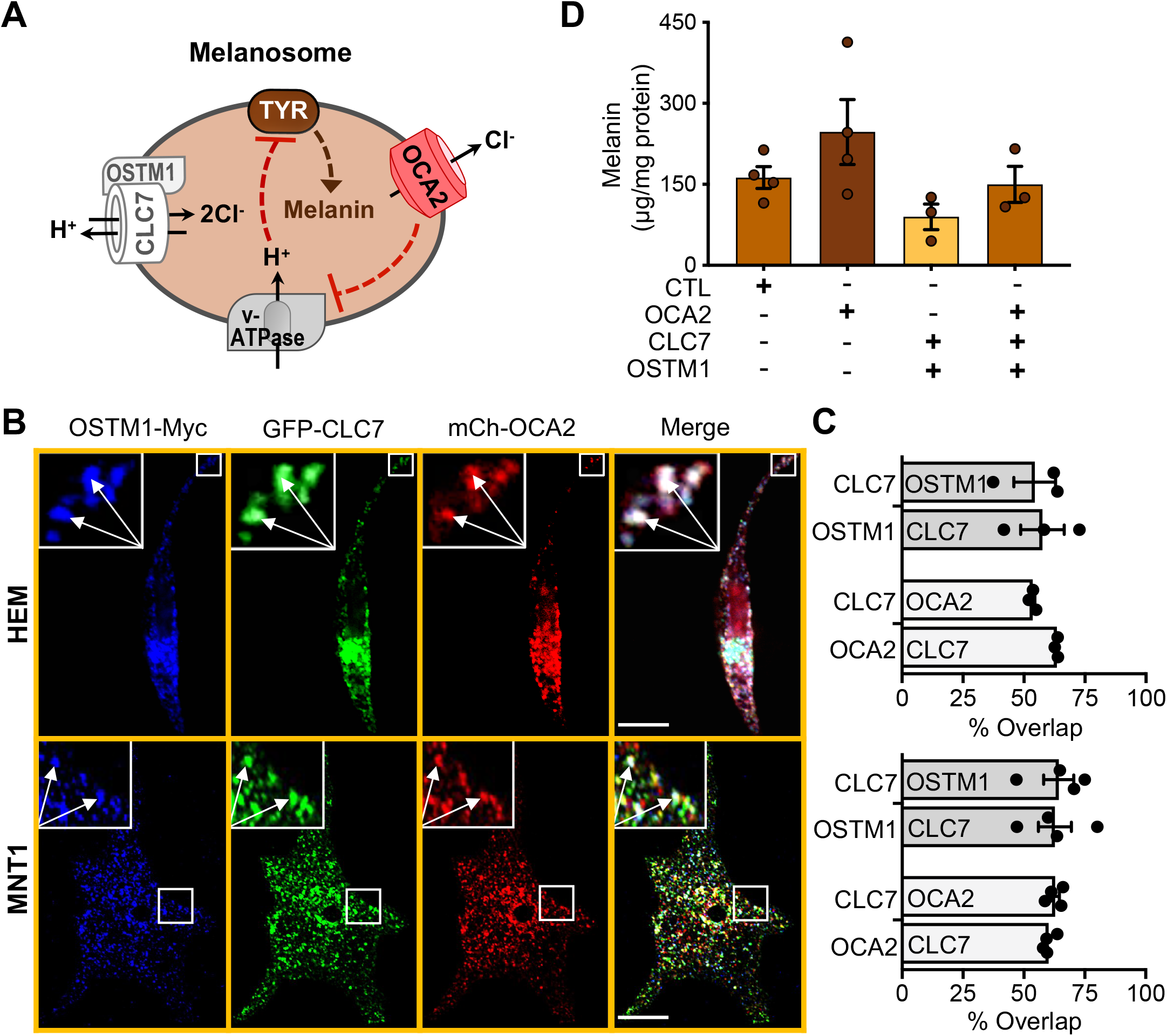
CLC7 colocalizes with melanosomal OCA2 human melanocytes. **A.** Schematic representation of OCA2-mediated regulation of melanosome pH and melanin synthesis. Cl^−^ efflux via OCA2 decreases vacuolar ATPase (V-ATPase) activity resulting in elevated tyrosinase (TYR) activity and melanin synthesis. **B.** Representative confocal fluorescence micrographs of HEMs and MNT1 cells transiently transfected with GFP-CLC7, OSTM1-Myc, visualized by anti-Myc (α-Myc) immunostaining, and mCh-OCA2. Inset: Magnified view of the area in white boxe, with arrows indicating organelles coexpressing GFP-CLC7, OSTM1-Myc, and mCh-OCA2. Scale bar: 10 μm. **C.** Quantification of images as in **B** shows overlap between overexpressed GFP-CLC7 and OSTM1-Myc or mCh-OCA2 in HEMs (top) and MNT1 cells (bottom). In both HEMs and MNT1 cells, ≥64% and ≥62% of total GFP-CLC7 localizes to organelles expressing OSTM1-Myc or mCh-OCA2, respectively. At least 60% of organelles with mCh-OCA2 in HEMs and MNT1 cells colocalize with GFP-CLC7. ± SEM; n ≥ 3 independent experiments. **D.** Total cellular melanin concentration in MNT1 cells transiently expressing for five days mCh-OCA2 alone or together with GFP-CLC7 and OSTM1-Myc. Compared to MNT1 cells expressing CLC7 and OSTM1, melanin content is increased 2.8-fold in cells expressing OCA2 alone (P = 0.0867) and 1.7-fold when OCA2 was expressed together with CLC7 and OSTM1 (P = 0.2163). P-values derived from unpaired two-sample t-tests. ± SEM; n ≥ 3 independent experiments.

### Melanosome pH measurements with the novel ratiometric pH sensor RpHiMEL

The melanosome lumen displays a wide range of pH values: early-stage melanosomes are acidic and become more neutral as they mature and accumulate melanin. To test if CLC7 regulates melanosome pH, we designed a ratiometric pH sensor, named RpHiMEL for <u>r</u>atiometric <u>pH i</u>ndicator for <u>me</u>lanosomes, and monitored the pH of individual melanosomes in live cells. In the absence of a known melanosomal targeting sequence, we used the loss-of-function OCA2 variant (OCA2-V443I) that localizes to melanosomes, but has negligible Cl^−^ conductance (6), to target RpHiMEL to melanosomes (**Fig. 4A**). We inserted the pH-sensitive fluorescent protein Nectarine (35) in the first intraluminal loop of OCA2-V443I, similar to the single-emission pH sensor MELOPS (5), and fused the pH-insensitive iRFP fluorescent protein to the cytosolic N-terminus of OCA2-V443I. The ratiometric nature of the indicator, together with real-time tracking of melanosomes expressing RpHiMEL, allowed us to accurately measure the pH of individual melanosomes, including more acidic ones that exhibit very low Nectarine fluorescence.

**Figure 4.**
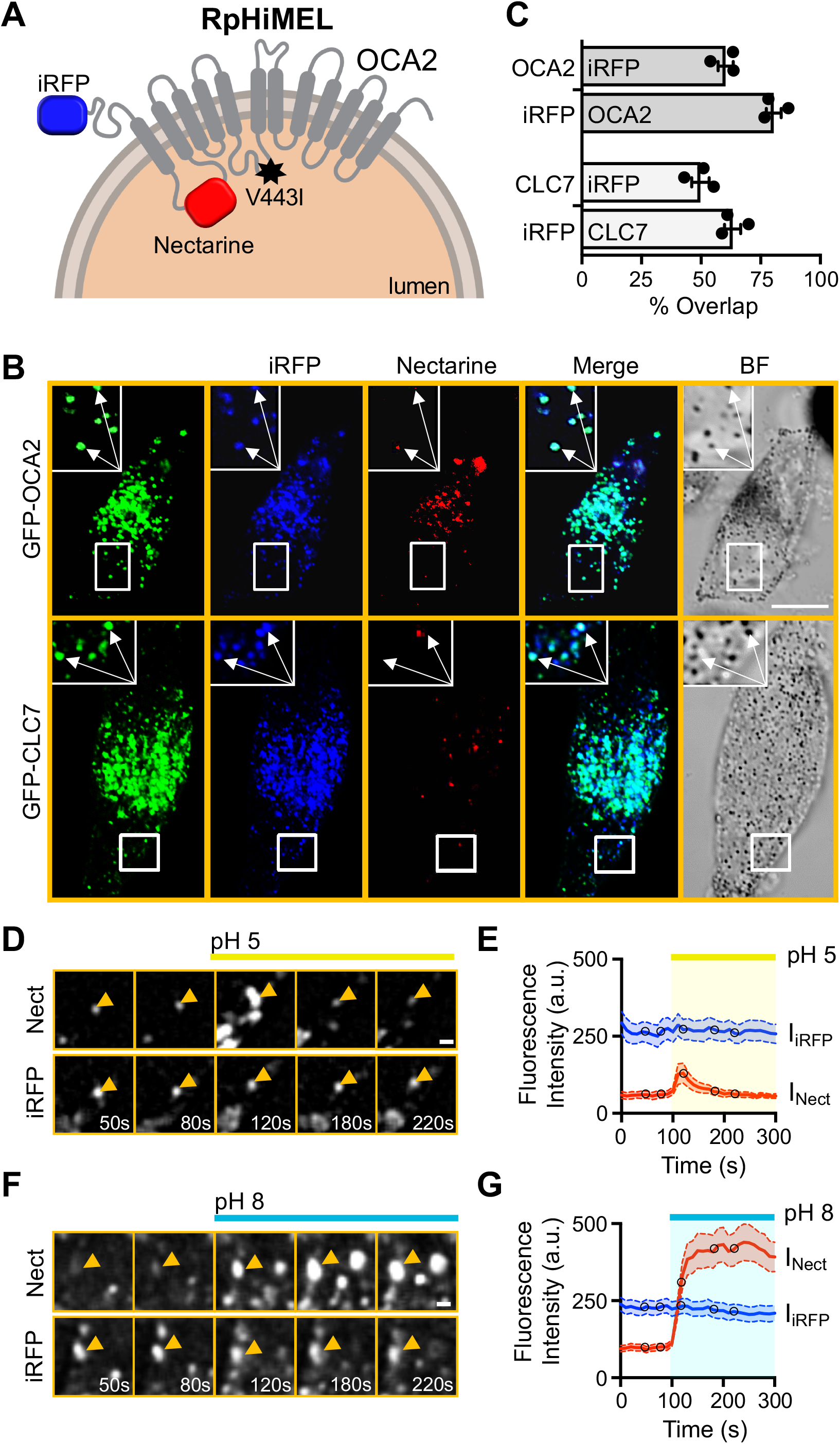
RpHimel is a novel fluorescent ratiometric indicator for melanosome pH. **A.** Schematic representation of the molecular design of RpHiMEL, a fluorescent ratiometric indicator for melanosome pH. RpHiMEL consists of the human melanosome-localized, loss-of-function OCA2-V443I mutant fused at its N-terminus to the pH-insensitive near-infrared fluorescent protein (iRFP) and containing the pH-sensitive Nectarine fluorescent protein in its first intraluminal loop. **B.** MNT1 human melanoma cells transiently expressing RpHiMEL and GFP-OCA2 (top) or GFP-CLC7 (bottom). Inset: Magnified view of boxed areas (white). Arrows indicate organelles coexpressing RpHiMEL and OCA2 or CLC7. Scale bar: 10 μm. **C.** Quantification of images as in **B** for reciprocal overlap between the iRFP of RpHiMEL and GFP-OCA2 (top) or GFP-CLC7 (bottom) in MNT1 cells. RpHiMEL colocalizes with 60% and 50% of all OCA2- or CLC7-expressing organelles, respectively. ± SEM; n = 3 independent experiments. **D & F.** Imaging of individual melanosomes in MNT1 cells expressing NT shRNA and RpHiMEL. Sequential images were acquired for the pH-sensitive Nectarine and the pH-insensitive iRFP before and after equilibration with pH 5 (**D**) or pH 8 (**F**) buffers. Arrowhead indicates the same organelle expressing RpHiMEL at different time points during the experiment. Yellow and blue bars represent the addition of pH 5 or pH 8 buffer plus ionophores, respectively. Scale bar: 2 μm. **E & G.** Average fluorescence intensity of Nectarine (I_Nect_) and iRFP (I_iRFP_) of RpHiMEL-expressing melanosomes as a function of time. Baseline images were taken for 100 s prior to equilibration with pH 5 (**E**) or pH 8 (**G**) buffer plus ionophores. At pH 5, I_Nect_ shows a small transient increase, while I_iRFP_ is unchanged. At pH 8, I_Nect_ shows a large and persistent increase, while I_iRFP_ is unchanged. Circles represent the time points at which the images in **D** and **F** were acquired. Yellow and blue bar represents the addition of pH 5 or pH 8 buffer plus ionophores, respectively. ± 95% confidence interval. Data from 60–80 melanosomes per pH value from n = 3 independent experiments. a.u., arbitrary units.

We validated the melanosomal localization of RpHiMEL in MNT1 cells and as expected, RpHiMEL overlaps with GFP-CLC7 and GFP-OCA2 (50% and 80%, respectively), demonstrating that RpHiMEL is properly trafficked to melanosomes containing both CLC7 and OCA2 (**Fig. 4B, 4C**). To determine the pH of individual melanosomes, we acquired time-lapse fluorescence images of MNT1 cells expressing RpHiMEL at 10 s intervals before and after the addition of pH buffers together with ionophores. Individual melanosomes were manually tracked and their iRFP and Nectarine fluorescence emissions were measured as a function of time (**Fig. 4D-4G**). In response to pH 5 buffer, we observed a small and transient increase in Nectarine fluorescence (I_Nect_), most likely due to the pH equilibration between the cytosol and melanosomes, but no change in iRFP fluorescence (I_iRFP_; **Fig. 4D, 4E**). Following the addition of pH 8 buffer, however, we observed a large and sustained increase in I_Nect_, but no change in I_iRFP_ (**Fig. 4F, 4G**). These results indicate that the RpHIMEL ratio I_Nect_/I_iRFP_ can be used to measure pH values between 5 and 8.

### Melanocytes with reduced CLC7 expression have less acidic melanosomes

To test whether CLC7 regulates melanosomal pH, we expressed RpHiMEL in MNT1 cells stably expressing NT or CLC7-targeting shRNA. Individual melanosomes from control NT shRNA cells displayed a large and persistent increase in I_Nect_, but no change in I_iRFP_ when equilibrated with pH 8 buffer (**Fig. 5A, 5B**). Melanosomes with reduced CLC7 expression, however, exhibited a smaller increase in I_Nect_ in response to pH 8 and no change in I_iRFP_ (**Fig. 5C, 5D**). This smaller increase in I_Nect_ appeared to be the result of an elevated I_Nect_ baseline fluorescence rather than a lower maximal value. At baseline (t < 100 s), I_Nect_ ~ I_iRFP_ in melanosomes from CLC7 shRNA cells, but not in NT shRNA cells, where I_Nect_ < I_iRFP_ (**Fig. 5B, 5D**).

**Figure 5.**
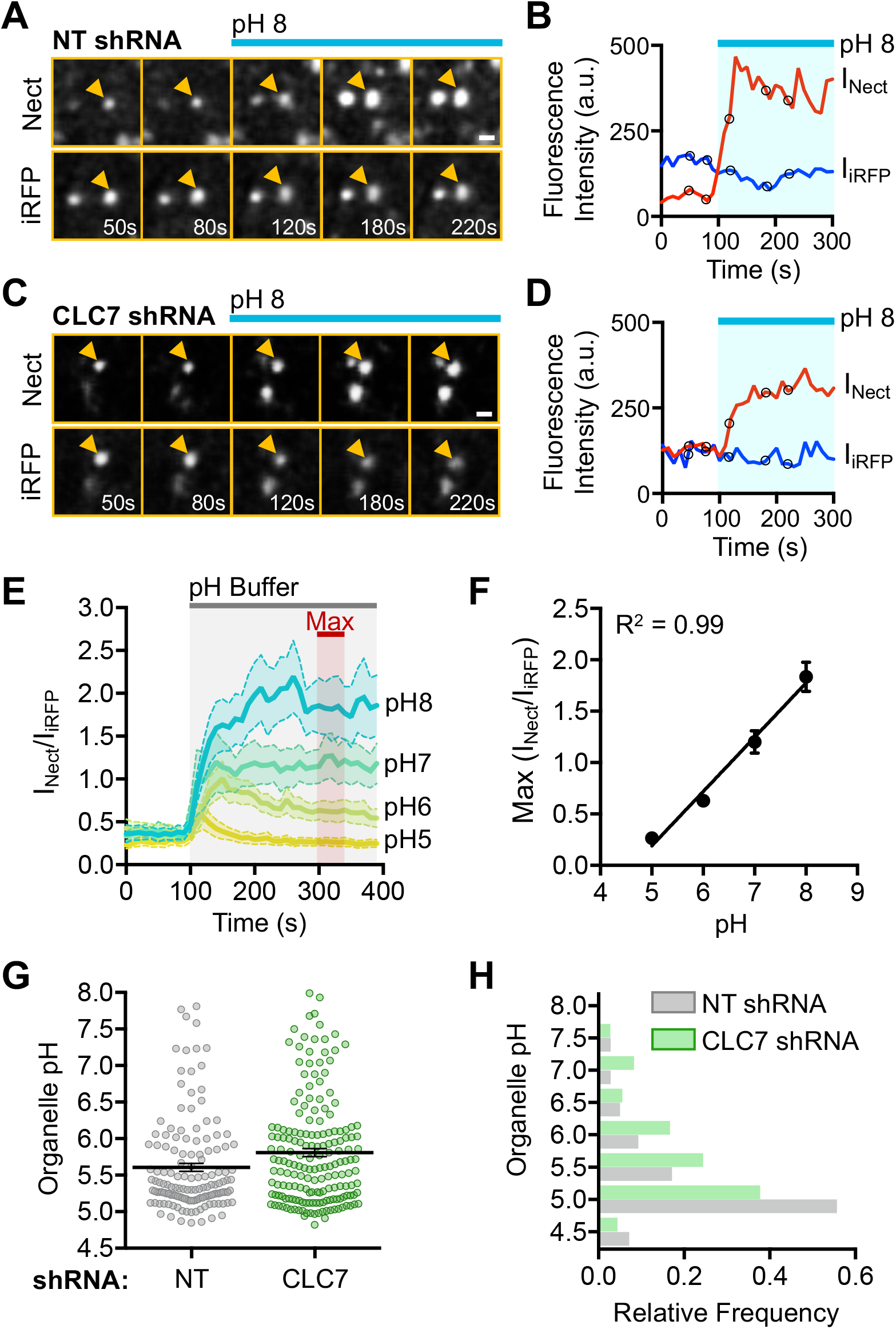
CLC7 regulates melanosome pH. **A & C.** pH-sensitive Nectarine and pH-insensitive iRFP channels from RpHiMEL imaging of individual melanosomes in MNT1 cells stably expressing NT shRNA (**A**) or CLC7 shRNA (**C**) before and after equilibration with pH 8 buffer. Arrowhead indicates the same organelle expressing RpHiMEL. Blue bar represents the addition of pH 8 buffer plus ionophores. Scale bar: 2 μm. **B & D.** Fluorescence intensity of iRFP (I_iRFP_) and Nectarine (I_Nect_) as a function of time for the melanosomes indicated by the arrowhead in **A** and **C**. Baseline images were taken for 100 s prior to equilibration with pH 8 buffer plus ionophores to obtain the maximal I_Nect_ fluorescence. Circles represent the time points at which the images in **A** and **C** were acquired. Blue bar represents the addition of pH 8 buffer plus ionophores. a.u., arbitrary units. **E.** Changes in the average RpHiMEL fluorescence intensity ratio (I_Nect_/I_iRFP_) of melanosomes from MNT1 cells stably expressing NT shRNA. After recording baseline fluorescence for 100 s, melanosomes were equilibrated with pH 5–8 buffers plus ionophores (grey bar). For each melanosome, maximal I_Nect_/I_iRFP_ was defined as the average I_Nect_/I_iRFP_ from t = 300–350 s (Max; red bar). Data represent 60–80 melanosomes per pH value from n = 3 independent experiments. ± 95% confidence interval. **F.** RpHiMEL calibration curve represented as a function of pH and obtained as the average maximal I_Nect_/I_iRFP_ measured in **E.** ± SEM. I_Nect_/I_iRFP_ is linear for pH values between 5 and 8 (R^2^ = 0.99; Y = 0.5278X – 2.448). **G.** pH values of individual melanosomes from MNT1 cells stably expressing NT shRNA (grey circles) or CLC7 shRNA (green circles). The average melanosomal pH of control NT and CLC7 shRNA cells is 5.61 ± 0.06 and 5.81 ± 0.06, respectively (P = 0.0104, Unpaired two-sample t-test). Black lines represent mean ± SEM. Data represent 140–180 organelles from n = 5 independent experiments. **H.** Frequency distribution of melanosome pH in MNT1 cells stably expressing NT shRNA (grey bar) or CLC7 shRNA (green bar). Melanosome pH in CLC7 shRNA cells is shifted towards higher pH values compared to control NT cells (P = 0.0014, Kolmogorov-Smirnov test).

To generate a pH calibration curve for RpHIMEL, we imaged MNT1 cells expressing RpHiMEL before and after addition of pH 5, 6, 7, or 8 buffers and calculated the average I_Nect_ /I_iRFP_ ratio of individual melanosomes from multiple cells as a function of time (**Fig. 5E**). The average I_Nect_ /I_iRFP_ ratio measured between 300-350 s for each pH buffer were fitted to obtain a standard curve (**Fig. 5F**). We then determined the pH of melanosomes from MNT1 cells stably expressing NT or CLC7-targeting shRNA and found that cells with reduced CLC7 expression have a higher average pH (5.81 ± 0.06) compared to control cells (5.61 ± 0.06) (**Fig 5G**). In addition, the relative frequency distribution of the pH values of individual melanosomes shows a shift to higher pH for melanosomes with reduced CLC7 expression, suggesting that CLC7 regulates melanin synthesis by acidifying melanosomes and, consequently, decreasing the activity of the melanogenic enzyme tyrosinase (TYR) (**Fig 5H**).

### Melanocytes expressing the CLC7-Y715C variant have large acidic vacuoles and reduced pigmentation

Recently, a *de novo* mutation in CLC7 (CLC7-Y715C) was identified in two unrelated young patients presenting a complex developmental phenotype, including albinism (18). At the cellular level, the Y715C variant of CLC7 has increased transporter activity, resulting in more acidic lysosomes and markedly enlarged intracellular vacuoles in patient fibroblasts (18). This gain-of-function variant is dominant, causing the same phenotype when expressed in wild-type fibroblasts (18). We sought to determine if the Y715C variant of CLC7 affects the ability of human melanocytes to produce melanin. MNT1 cells expressing the Y715C variant exhibited drastically different morphology with multiple large cytoplasmic vacuoles (**Fig. 6A, 6B**) similar to the morphology phenotype of CLC7-Y715C patient fibroblasts (18). The vacuoles were visible due to exclusion of expressed cytosolic mCh; nevertheless, to quantify the phenotype of the CLC7-Y715C expressing cells, we tested different fluorescent probes for accumulation in or association with the membrane delimiting the vacuoles. Interestingly, the ratiometric pH indicator LysoSensor DND-160 accumulated in the same vacuoles that exclude cytosolic mCh, allowing us to measure the size of organelles, and their relative acidity (**Fig. 6B, 6C**).

**Figure 6.**
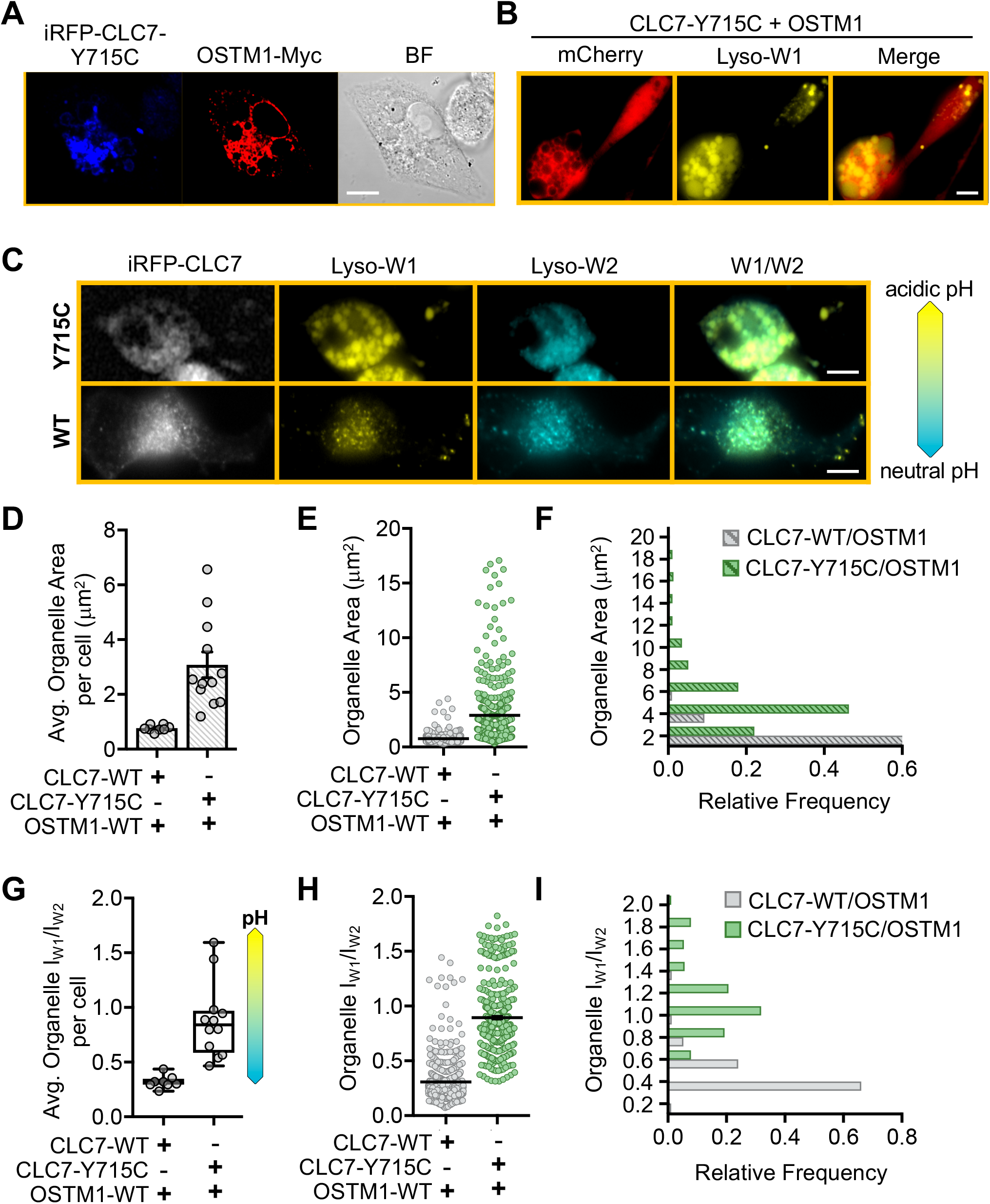
Melanocytes expressing the CLC7-Y715C mutant have enlarged acidic vacuoles. **A.** MNT1 cells expressing iRFP-CLC7-Y715C and OSTM1-Myc have enlarged cytosolic vacuoles, as observed by bright field (BF) microscopy. OSTM1-Myc was detected by anti-Myc immunostaining. Scale bar: 10 μm. **B.** Wide-field micrographs of MNT1 cells expressing mCh, iRFP-CLC7-Y715C, and OSTM1-Myc and incubated with LysoSensor show that the enlarged cytosolic vacuoles that exclude mCh accumulate LysoSensor, visualized using its yellow fluorescent emission (Lyso-W1). Scale bar: 10 μm. **C.** Representative widefield fluorescence images of LysoSensor-stained MNT1 cells transiently coexpressing OSTM1-Myc with either iRFP-CLC7-Y715C (top) or iRFP-CLC7-WT (bottom). LysoSensor dual fluorescence emission Lyso-W1 (yellow) and Lyso-W2 (blue), and the ratio W1/W2 are shown in pseudocolor, with more yellow W1/W2 indicating increased organelle acidity, as shown by the pH scale. Scale bar: 10 μm. **D.** Average cellular organelle area (in μm^2^) obtained as individual LysoSensor-stained organelles area the averaged for individual MNT1 cells expressing the indicated proteins. The average organelle area per MNT1 cell transiently expressing OSTM1-Myc with iRFP-CLC7-WT or iRFP-CLC7-Y715C is (0.77 ± 0.04) μm^2^ and (3.08 ± 0.47) μm^2^, respectively. (P < 0.0001, Mann-Whitney test). Bar graphs represent mean ± SEM. **E.** Surface area of individual LysoSensor-stained organelles from MNT1 cells expressing OSTM1-Myc together with iRFP-CLC7-WT (grey circles) or iRFP-CLC7-Y715C (green circles). The average organelle area in cells with CLC7-Y715C is 3.8-fold higher compared to CLC7-WT cells (P < 0.0001, Mann-Whitney test), indicating that CLC7-Y715C overexpression results in increased organelle size. Black lines represent mean ± SEM. **F.** Frequency distribution of organelle area from values obtained in **E**. CLC7-Y715C expression shifts the organelle area distribution to higher values, indicating increased organelle size in CLC7-Y715C cells compared to CLC7-WT cells (P < 0.0001, Kolmogorov-Smirnov test). **G.** Average cellular ratio of the two emission intensities of LysoSensor (I_W1_/I_W2_) obtained as the average of LysoSensor-stained organelles from individual MNT1 cells expressing the indicated proteins. The average organelle IW1/IW2 per MNT1 cell transiently expressing OSTM1-Myc with iRFP-CLC7-WT or iRFP-CLC7-Y715C is 0.32 ± 0.02 and 0.88 ± 0.10, respectively (± SEM; P < 0.0001, Mann-Whitney test). A higher I_W1_/I_W2_ indicates increased organelle acidity as shown by the pH scale (right). Box plots show the data medians, 25^th^ and 75^th^ percentiles, and extrema. **H.** I_W1_/I_W2_ values of individual LysoSensor-containing organelles from MNT1 cells overexpressing OSTM1-Myc with iRFP-CLC7-WT (grey circles) or iRFP-CLC7-Y715C (green circles). The average organelle I_W1_/I_W2_ in cells with CLC7-Y715C is increased 2.8-fold compared to CLC7-WT cells, indicating that CLC7-Y715C increases organelle acidity (P < 0.0001, Mann-Whitney test). Black lines represent mean ± SEM. **I.** Frequency distribution of organelle IW1/IW2 from values obtained in **H**. CLC7-Y715C expression shifts the organelle I_W1_/I_W2_ distribution to higher values, indicating increased organelle acidity in CLC7-Y715C cells compared to CLC7-WT cells (P < 0.0001, Kolmogorov-Smirnov test). Organelle I_W1_/I_W2_ measurements in **G-I** were paired with organelle area measurements in **D-F**. ≥ 367 organelles/cell from ≥ 8 cells were analyzed; n = 5 independent experiments.

In MNT1 cells expressing CLC7-Y715C and OSTM1, LysoSensor accumulated in cytosolic organelles that were larger and appeared more acidic (higher yellow emission) than in cells expressing WT CLC7 and OSTM1 (**Fig. 6C**). The cellular average of the cross-sectional area of LysoSensor-positive organelles in MNT1 cells expressing the Y715C variant was approximately 3-fold larger compared to WT CLC7-expressing cells (**Fig. 6D**). The area of individual organelles detected on both yellow (W1) and blue (W2) emission wavelengths of LysoSensor was also higher and covered a wider range for MNT1 cells expressing CLC7-Y715C compared to WT CLC7 (**Figs. 6E, 6F**). For the same LysoSensor-containing organelles used for area measurements, we calculated the ratio of the yellow and blue emission intensities (IW1/IW2), as an indicator of organelle pH. The cellular IW1/IW2 average was ~ 2-fold higher for the larger organelles detected in CLC7-Y715C-expressing cells than in those expressing WT CLC7 (**Fig. 6G**). The individual LysoSensor-containing compartments were also significantly more acidic in cells expressing the Y715C variant, compared to WT CLC7 (**Fig. 6H, 6I**), suggesting that expression of CLC7-Y715C causes not only more acidic lysosomes (18), but also the formation of acidic vacuoles.

### OCA2 restores pigmentation in hypopigmented melanocytes expressing the CLC7-Y715C variant

In addition to the change in morphology, MNT1 cells expressing CLC7-Y715C and OSTM1 have reduced melanin content (**Fig. 7A**), possibly due to more acidic melanosomes with reduced TYR activity and lower melanin synthesis rates. Because the majority of melanosomes expressing CLC7 and OSTM1 also express OCA2 (**Fig. 3B, 3C**), we tested if coexpression of OCA2, a positive regulator of melanosomal pH (6), rescues the hypopigmentation phenotype of CLC7-Y715C. Indeed, ectopic expression of OCA2 in cells expressing CLC7-Y715C restored pigmentation to WT levels (**Fig. 7A**). We wondered if ectopic expression of OCA2 in melanocytes could restore not only pigmentation, but also prevent the morphological phenotype. Using as a control the OCA2-V443I mutant, which has negligible effects on melanosomal pH (6), we showed that either WT or V443I OCA2 ectopically expressed in MNT1 cells remained partially colocalized with CLC7-Y715C and OSTM1 (**Fig. S4A**). In addition, expression of either WT or V443I OCA2 alone did not change the area of LysoSensor-positive organelles of these cells (**Fig. S4B, S4C**), while only WT OCA2 neutralized the pH of the compartments that accumulate LysoSensor (**Fig. S4D, S4E**).

**Figure 7.**
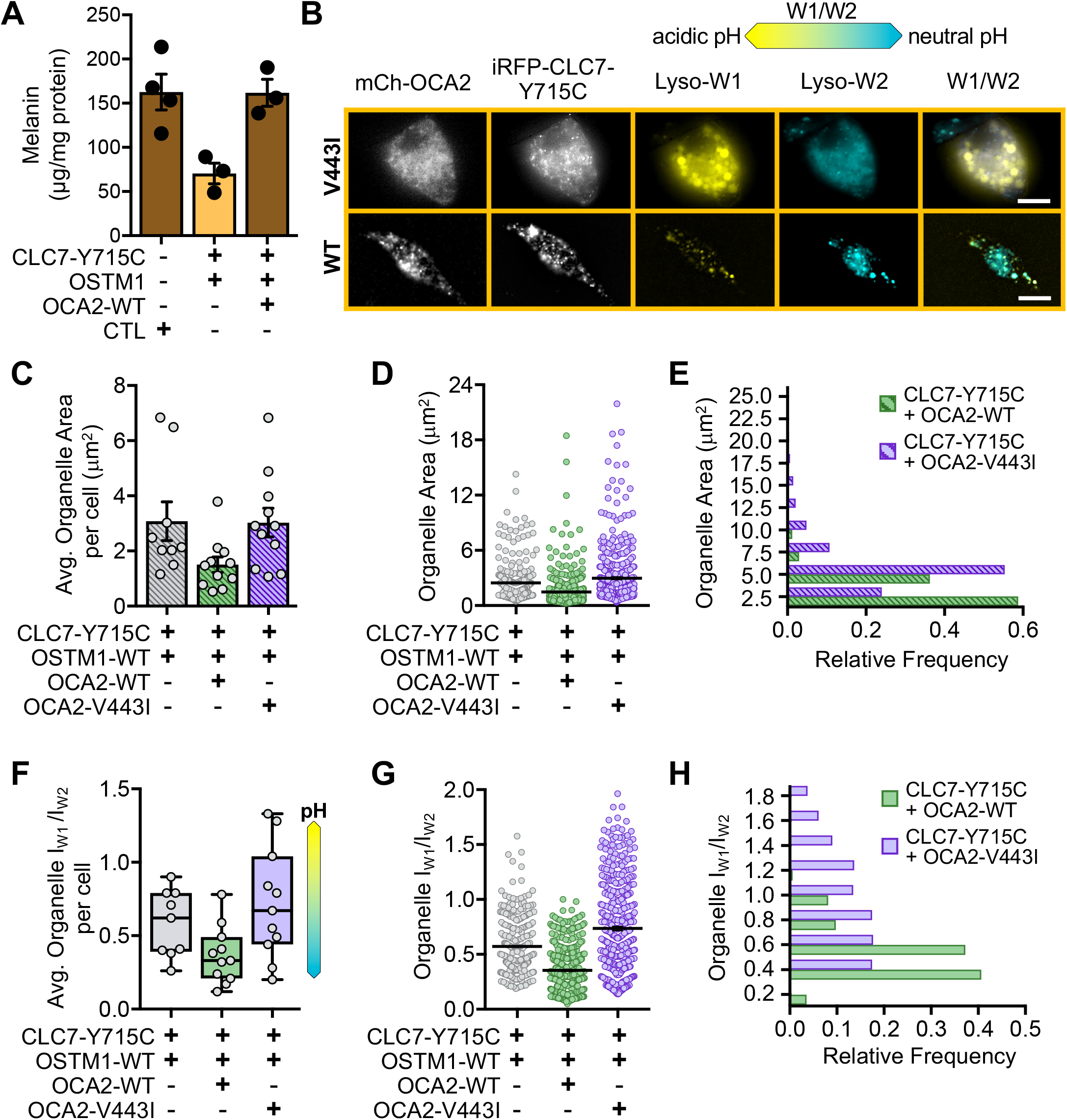
The phenotype of MNT1 human melanoma cells expressing gain-of-function CLC7-Y715C mutant is restored by ectopic expression of OCA2. **A.** Total cellular melanin concentration of MNT1 cells transiently expressing iRFP-CLC7-Y715C and OSTM1-Myc (orange bar) is decreased 2.3-fold compared to mock-transfected control cells (left brown bar, P = 0.0166, One-way ANOVA with Tukey’s multiple comparisons test), and is restored to control levels by ectopic expression of WT mCh-OCA2 (right brown bar; P = 0.9994, One-way ANOVA with Tukey’s multiple comparisons test). Bars represent mean values ± SEM; n ≥ 3 independent experiments. **B.** Representative widefield fluorescence images of LysoSensor-stained MNT1 cells transiently expressing iRFP-CLC7-Y715C and OSTM1-Myc with either mCh-OCA2-V443I (top) or mCh-OCA2-WT (bottom). LysoSensor dual fluorescence emission Lyso-W1 (yellow) and Lyso-W2 (blue) and the ratio W1/W2 are shown in pseudocolor, with more yellow W1/W2 indicating increased organelle acidity, as shown by the pH scale. Scale bar: 10 μm. **C.** Average cellular organelle area (in μm^2^) obtained as the average of LysoSensor-stained organelles from individual MNT1 cells expressing the proteins indicated below. The average organelle area per MNT1 cell expressing iRFP-CLC7-Y715C and OSTM1-Myc alone (3.08 ± 0.70 μm^2^) was not significantly changed by coexpression with mCherry-OCA2-V443I (3.03 ± 0.52 μm^2^), while coexpression of mCherry-OCA2-WT decreased the average organelle area to 1.50 ± 0.28 μm^2^ (P = 0.0879, One-way ANOVA with Tukey’s multiple comparisons test). Each dot represents the average organelle area of one cell. Bars represent the mean of all cells per condition ± SEM. **D.** Area of individual LysoSensor-stained organelles from MNT1 cells overexpressing iRFP-CLC7-Y715C and OSTM1-Myc (grey circles) with mCherry-OCA2-WT (green circles) or mCherry-OCA2-V443I (purple circles), as indicated. Compared to control cells expressing CLC7-Y715C and OSTM1 alone or with OCA2-V443I, the average organelle area is decreased by ≥ 2.0-fold when OCA2-WT is coexpressed (P < 0.0001, Kruskal-Wallis test with Dunn’s multiple comparisons test). Black lines represent the mean ± SEM. **E.** Frequency distribution of organelle area values obtained in **D**. Compared to control cells expressing CLC7-Y715C and OSTM1 with OCA2-V443I, coexpression of OCA2-WT shifts the organelle area distribution to lower values, indicating reduced organelle size (P < 0.0001, Kolmogorov-Smirnov test). **F.** Average cellular ratio (I_W1_/I_W2_) of the fluorescence intensity of Lyso-W1 (I_W1_) and Lyso-W2 (I_W2_) of individual LysoSensor-stained organelles from individual MNT1 cells expressing the indicated proteins. The average organelle I_W1_/I_W2_ per cell in MNT1 overexpressing iRFP-CLC7-Y715C and OSTM1-Myc alone or together with mCherry-OCA2-V443I were similar, 0.64 ± 0.07 and 0.72 ± 0.11, respectively, while coexpression of mCh-OCA2-WT resulted in lower IW1/IW2 values (0.37 ± 0.06), indicating less acidic organelles. (P = 0.0180; One-way ANOVA with Tukey’s multiple comparisons test). Dots represent individual cells, box plots show the data medians, 25^th^ and 75^th^ percentiles, and extrema. **G.** I_W1_/I_W2_ values of individual LysoSensor-stained organelles from MNT1 cells overexpressing iRFP-CLC7-Y715C and OSTM1-Myc (grey circles) with mCherry-OCA2-WT (green circles) or mCherry-OCA2-V443I (purple circles). Compared to control cells expressing CLC7-Y715C and OSTM1 with or without OCA2-V443I, the average organelle I_W1_/I_W2_ is decreased by ≥ 1.6-fold in cells rescued with OCA2-WT (P < 0.0001; Kruskal-Wallis test with Dunn’s multiple comparisons test). Mean ± SEM **H.** Frequency distribution of organelle IW1/IW2 values shown in **G**. Compared to control cells expressing CLC7-Y715C and OSTM1 with OCA2-V443I, OCA2-WT coexpression shifts the organelle IW1/IW2 distribution to lower values, indicating reduced organelle acidity (P < 0.0001, Kolmogorov-Smirnov test). Organelle I_W1_/I_W2_ measurements in **F-H** were paired with organelle area measurements in **C-E**. Area and I_W1_/I_W2_ measurements were acquired from ≥ 253 organelles per cell from at least 9 cells; n = 3 independent experiments.

When we expressed V443I or WT OCA2 together with CLC7-Y715C and OSTM1 and used LysoSensor (**Fig. 7B**), it became apparent that on average cells expressing OCA2 WT, but not V443I, lacked the large acidic vacuoles typical for the CLC7-Y715C variant. To quantify the effect of OCA2 coexpression, we measured the cellular average area of LysoSensor-positive organelles, which showed significantly smaller acidic organelles in cells expressing CLC7-Y715C and OCA2-WT, while OCA2-V443I expression had no effect on the cellular average of acidic organelle size (**Fig. 7C**). The average and distribution of organelle area was also significantly decreased and shifted, respectively, in MNT1 cells expressing CLC7-Y715C and OCA2-WT, but not OCA2-V443I (**Fig. 7D, 7E**). Interestingly, not only was the size of the LysoSensor compartments reduced by WT-OCA2 expression, but also their pH became less acidic, as measured by the LysoSensor emission ratio I_W1_/I_W2_ (**Fig. 7F-7H**). Likewise, mCh-OCA2, but not mCh alone, rescued the large and acidic pH of organelles from cells coexpressing mCh-OCA2 and CLC7-Y715C (**Fig. S4F-S4J**).

### OCA2 expression restores the cellular phenotype of the CLC7-Y715C variant in non-melanocytic cells

Because ectopically expressed OCA2 localizes to lysosomes in non-melanocytic cells, similar to CLC7 and OSTM1 (>70% overlap with LAMP1-positive organelles in HeLa cells, **Fig. S3C, S3D**) (6) (36), we investigated whether OCA2 could also rescue the morphology phenotype caused by the CLC7-Y715 mutant in non-melanocytic cells. HeLa cells expressing CLC7-Y715 and OSTM1 (**Fig. S5A**) showed a phenotype similar to MNT1 cells (**Fig. 6A**) and patient fibroblasts (18), and the resulting vacuoles also accumulated LysoSensor (**Fig. S5B, S5C**). HeLa cells expressing CLC7-Y715C had a cellular average organelle area that was approximately 8-fold higher than cells expressing CLC7-WT (**Fig. S5D**), which was similar to the difference in the area and size distribution of individual organelles between each cell population (**Fig. S5E, S5F**). In addition, LysoSensor-positive organelles in HeLa cells expressing the CLC7-Y715C variant were significantly more acidic compared to HeLa cells expressing CLC7-WT and OSTM1 (**Fig. S5G-S5I**). Ectopically expressed OCA2-WT or OCA2-V443I partially colocalized with the CLC7-Y715C variant and with OSTM1 in HeLa cells (**Fig. S6A**). The expression of only OCA2-WT or -V443I did not alter the size of acidic organelles (**Fig. S6B, S6C**), while OCA2-WT, but not V443I shifted the pH of LysoSensor-positive organelles to more neutral values, as expected (**Fig. S5D, S5E**) (6). We next coexpressed OCA2-WT with CLC7-Y715C and OSTM1 and observed that, unlike the expression of OCA2-V443I, the phenotype of the cells appears restored with respect to the size and acidity of the organelles containing LysoSensor (**Fig. 8A**). Indeed, the average organelle area per HeLa cell expressing CLC7-Y715C and OSTM1 together with OCA2-WT, but not OCA2-V443I, was reduced more than 4-fold compared to HeLa cells expressing CLC7-Y715C alone (**Fig. 8B**). The area and size distribution of individual organelles was also significantly reduced and shifted, respectively (**Fig. 8C, 8D**). Similar to the phenotype in MNT1 cells, HeLa cells expressing CLC7-Y715C and OSTM1 with OCA2-WT, but not OCA2-V443I, had fewer LysoSensor-positive organelles with an acidic pH (**Fig. 8E-8G**).

**Figure 8.**
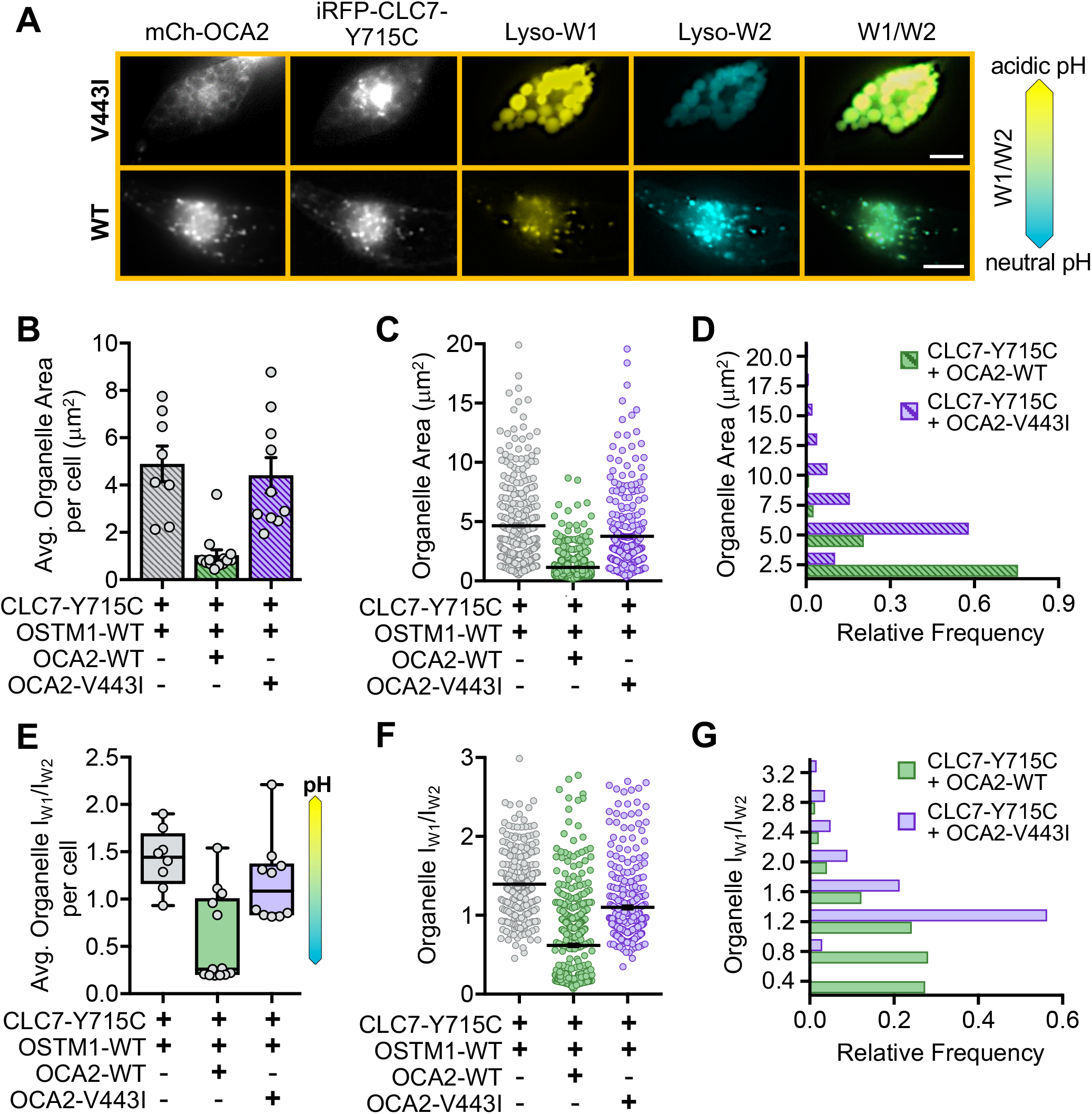
OCA2 expression restores vacuolar phenotype and pH in HeLa cells expressing CLC7-Y715C. **A.** Representative widefield fluorescence images of LysoSensor-stained HeLa cells expressing iRFP-CLC7-Y715C and OSTM1-Myc with mCh-OCA2-V443I (top) or mCh-OCA2-WT (bottom). LysoSensor dual fluorescence emission Lyso-W1 (yellow) and Lyso-W2 (blue) and the ratio W1/W2 are in pseudocolor, with more yellow W1/W2 indicating higher acidity as shown by the pH scale. Scale bar: 10 μm. **B.** Average organelle area (in μm^2^) per cell obtained as the average of LysoSensor-stained organelles from individual HeLa cells expressing the indicated proteins. The average organelle area per HeLa cell overexpressing iRFP-CLC7-Y715C and OSTM1-Myc with mCherry-OCA2-WT or mCherry-OCA2-V443I is 1.04 ± 0.23 μm^2^ and 4.41 ± 0.77 μm^2^, respectively (± SEM; P = 0.0008, Kruskal-Wallis test with Dunn’s multiple comparisons test). Each dot represents one cell. Bars represent the mean of all cells per condition ± SEM. **C.** Area values of individual LysoSensor-stained organelles from HeLa cells overexpressing iRFP-CLC7-Y715C and OSTM1-Myc (grey circles) with mCh-OCA2-WT (green circles) or mCh-OCA2-V443I (purple circles). Compared to control cells expressing CLC7-Y715C and OSTM1 with or without OCA2-V443I, the average organelle area is decreased by ≥ 3.3-fold in cells rescued with OCA2-WT (P < 0.0001, Kruskal-Wallis test with Dunn’s multiple comparisons test). Black lines represent mean ± SEM. **D.** Frequency distribution of organelle area values in **C**. Compared to control cells expressing CLC7-Y715C and OSTM1 with OCA2-V443I, OCA2-WT coexpression shifts the organelle area distribution to lower values, indicating that functional OCA2 rescues organelle size (P < 0.0001, Kolmogorov-Smirnov test). **E.** Average cellular IW1/IW2 obtained as the average of LysoSensor-stained organelles from individual HeLa cells expressing the indicated proteins. The average organelle IW1/IW2 per HeLa cell overexpressing iRFP-CLC7-Y715C and OSTM1-Myc with mCh-OCA2-WT or mCh-OCA2-V443I is 0.56 ± 0.13 and 1.81 ± 0.14, respectively (± SEM; P = 0.0049, One-way ANOVA with Tukey’s multiple comparisons test). Dots represent individual cells. Box plots show the data medians, 25^th^ and 75^th^ percentiles, and extrema. **F.** I_W1_/I_W2_ values of individual LysoSensor-stained organelles from HeLa cells overexpressing iRFP-CLC7-Y715C and OSTM1-Myc (grey circles) with mCh-OCA2-WT (green circles) or mCh-OCA2-V443I (purple circles). Compared to control cells expressing CLC7-Y715C and OSTM1 with or without OCA2-V443I, the average organelle I_W1_/I_W2_ is ≥ 1.7-fold lower in cells rescued with OCA2-WT (P < 0.0001; Kruskal-Wallis test with Dunn’s multiple comparisons test). Black lines represent mean ± SEM. **G.** Frequency distribution of organelle I_W1_/I_W2_ values in **F**. Compared to control cells expressing CLC7-Y715C and OSTM1 with OCA2-V443I, OCA2-WT coexpression shifts the organelle IW1/IW2 distribution to lower values, indicating reduced organelle acidity (P < 0.0001, Kolmogorov-Smirnov test). Organelle I_W1_/I_W2_ measurements in **E-G** were paired with organelle area measurements in **B-D** acquired from ≥ 281 organelles from at least 8 cells per condition; n = 3 independent experiments.

## Discussion

In conclusion, our data show that in melanocytes, CLC7 and OSTM1, are expressed in both lysosomes and melanosomes. The fraction of CLC7 and OSTM1 localized to melanosomes acidifies the melanosome lumen, opposing the activity of the melanosomal Cl^−^ channel OCA2. Melanosomal pH dictates the activity of the melanogenic enzyme TYR, thereby regulating the rate of melanin synthesis and extent of pigmentation (**Fig. 9A**, middle). Melanocytes with reduced levels of CLC7 and OSTM1 will have more basic melanosomes, resulting in higher TYR activity and increased pigmentation (**Fig. 9A**, right). Melanosomes that express the gain-of-function CLC7-Y715C variant will have a more acidic pH and reduced melanin content (**Fig. 9A**, left). In non-melanocytic cells, CLC7 and OSTM1 acidify lysosomes (**Fig. 9B**, left), while the CLC7-Y715C variant causes lysosome-like vacuoles to become hyperacidic (18) (**Fig. 9B**, middle). The effect of the CLC7-Y715C variant can be rescued by expressing OCA2, which localizes to lysosomes in non-melanocytic cells and counteracts the effect of CLC7-Y715C on luminal pH (**Fig. 9B**, right).

**Figure 9.**
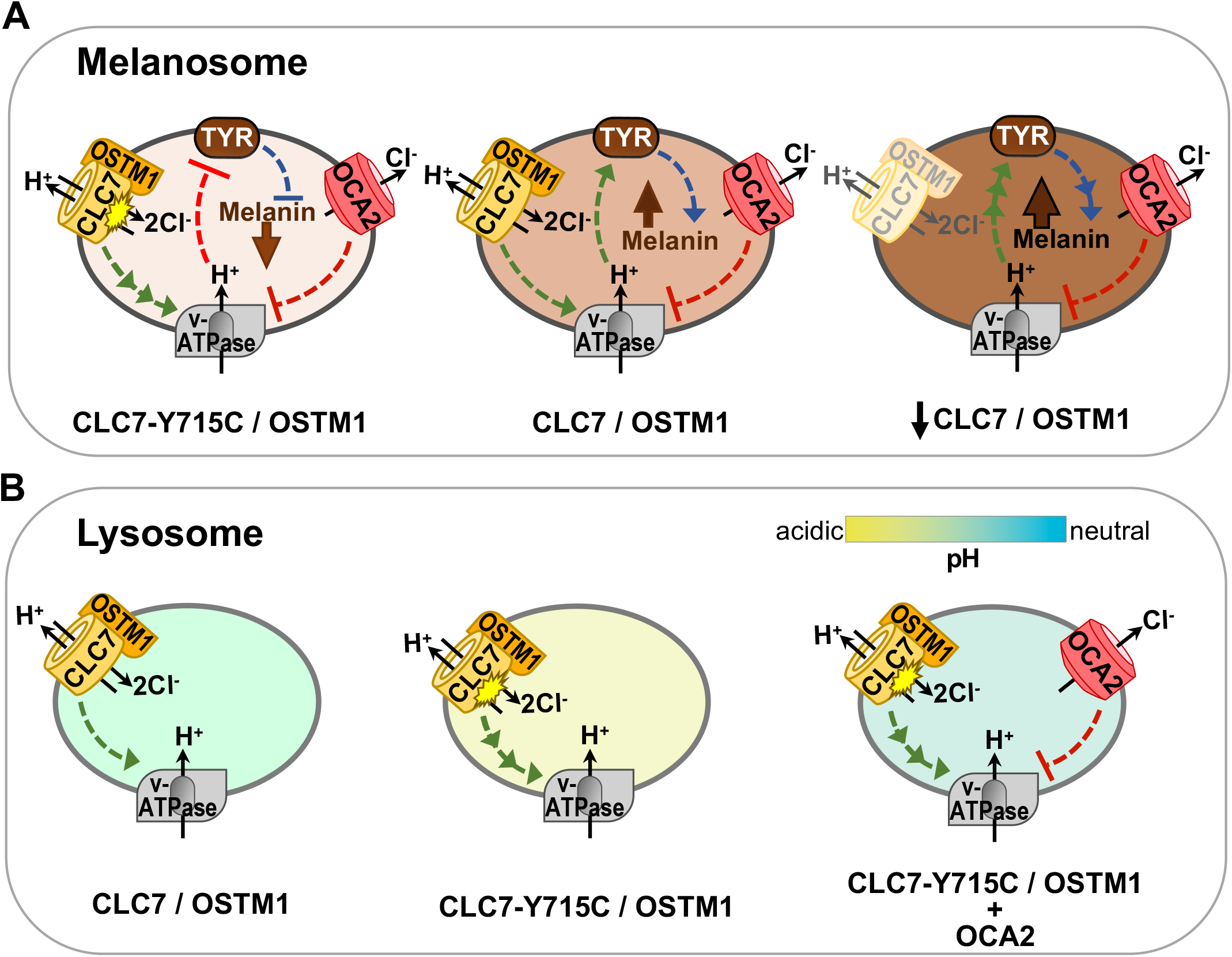
Proposed mechanism for melanosome pH regulation and melanogenesis by CLC7. **A.** Melanosomes are organelles in which melanin synthesis occurs, a process mediated by the main melanogenic enzyme TYR, whose catalytic activity is pH-dependent and optimal at near-neutral pH. The melanosomal lumen is acidified by the vesicular ATPase (V-ATPase). OCA2, a melanosomal chloride Cl^−^ channel, conducts Cl^−^ out of melanosomes, generating a surplus of luminal positive charges that decrease the activity of the electrogenic V-ATPase. Such action increases melanosome pH and TYR activity. CLC7/OSTM1 also reside in melanosomes and function together as a 2Cl^−^/1H^+^ antiporter, pumping Cl^−^ into the lumen, which increases the V-ATPase driving force and acidifies the melanosome lumen. Under physiological conditions (middle), CLC7/OSTM1 opposes the effect of OCA2 on V-ATPase and luminal pH. With loss of CLC7/OSTM1 expression (right), OCA2 activity leads to higher luminal pH and TYR activity that leads to increased melanin synthesis. Melanosomes expressing the gain-of-function CLC7-Y715C/OSTM1 mutant (left) have a more acidic luminal pH, which lowers TYR activity and decreases melanin levels. **B.** In lysosomes, CLC7/OSTM1 also increases the driving force of V-ATPase to acidify the lumen (left). Expression of the gain-of-function CLC7-Y715C/OSTM1 mutant (middle) further enhances the activity of V-ATPase, resulting in hyperacidified lysosomes and impaired organelle function, as observed in the patients carrying this mutation. The effect of CLC7-Y715C/OSTM1 on lysosomal pH can be restored if OCA2 is also expressed in these lysosomes (right) to counteract the increased V-ATPase activity and hyperacidic lysosome pH.

We conclude that CLC7/OSTM1 is a negative regulator of pigmentation in human skin melanocytes, as downregulation of CLC7 or its β-subunit OSTM1 results in increased pigmentation (**Fig. 1**), while overexpression of WT CLC7 or the gain-of function Y715C variant with OSTM1 causes reduced pigmentation (**Fig. 3D, Fig. 7A**). The pigmentation phenotype of human melanocytes only partially recapitulates the pigmentation phenotype of the CLC7^−/−^ and OSTM1^−/−^ mice, both of which show reduced pheomelanin levels (15, 17). In contrast, human melanocytes with CLC7 downregulation have increased pheomelanin, whereas OSTM1 downregulation leads to less pheomelanin (**Fig. 1E**), similar to the mouse hair phenotype. The unexpected difference in the direction of the pheomelanin changes in HEMs with downregulation of CLC7 compared to OSTM1 imply that OSTM1 might have cellular functions independent of CLC7, as previously suggested (16, 17). The difference between the effect of CLC7 downregulation on mouse and human pigmentation, while surprising, could be attributed to different mechanisms governing hair and skin pigmentation, or to the absence of CLC7 expression during early developmental stages in mice (14, 37). A similar difference between mouse hair and human skin pigmentation phenotypes was recently described for another transporter, MFSD12 (10, 38, 39): while MFSD12^−/−^ mice in agouti background are grey due to the absence of pheomelanin bands in the hair, human and mouse skin melanocytes have higher overall melanin levels (38).

Melanosomes and lysosomes tightly regulate their luminal pH, a critical factor for the luminal environment that regulates the activity of enzymes essential for the function of these organelles. Melanosomes also regulate their pH during each stage of maturation, with highly acidic pH observed in early melanosomes to almost neutral pH in mature melanosomes. Measuring melanosomal pH of individual melanosomes in live cells proved to be difficult, primarily due to pH variability between melanosomal stages and to melanin interfering with the fluorescence emission of pH-sensitive dyes. To address these problems, we designed a genetically-encoded ratiometric melanosomal pH indicator, RpHIMEL (**Fig. 4A**). Using RpHIMEL in live cells, we found that the melanosomes from cells expressing CLC7 shRNA have higher pH (**Fig. 5G, H**), unlike the pH of lysosomes obtained from CLC7^−/−^ mice, which is unchanged compared to controls (27). This difference in CLC7-mediated pH modulation for short term downregulation of CLC7 in melanocytes and deletion of CLC7 in mouse embryos, could be due to changes in the expression of other lysosomal channels and/or transporters during development in order to maintain the lysosomal pH, which is critical for cell function and survival. Similarly, when we express positive regulators of lysosomal pH, like OCA2 or SLC45A2, in heterologous cells (34), their effect on lysosomal pH can only be detected for a limited time (1-2 days), before the cells reestablish the acidic pH of the lysosomes.

Our finding that CLC7 partially localizes to a population of melanosomes expressing OCA2 (**Fig. 3B**) answers a key question raised by our previous study showing that OCA2 is a melanosomal Cl^−^ ion channel: by what mechanism is the melanosomal Cl^−^ replenished? As Cl^−^ needs to be transported against its electrochemical gradient, such a function could only be fulfilled by a melanosomal Cl^−^ transporter like CLC7. In addition to the dual regulation of melanosome pH by CLC7 and OCA2 and their colocalization, we also showed that ectopic expression of OCA2 compensates for the hyperacidic pH caused by the CLC7-Y715C variant (**Fig. 7F-7H**). Importantly, because OCA2 expressed in non-melanocytic cells localizes to lysosomes, its expression in HeLa cells restored the lysosomal defects and the changes in cell morphology caused by the CLC7-Y715C variant (**Fig. 8**), suggesting that OCA2 could rescue the pathogenic phenotypes observed in CLC7-Y715C patient cells.

In conclusion, our data uncovers a novel mechanism by which CLC7 regulates human skin pigmentation and identifies OCA2 as a tool to rescue the effects of CLC7 activating mutations in melanocytic and non-melanocytic cells.

## Materials and Methods

### Cell culture, transfection, and lentiviral transduction

Cell culture reagents were purchased from Thermo Fisher Scientific (Fisher Scientific, Waltham, MA) unless otherwise specified.

#### Normal primary human epidermal melanocytes (HEMs)

derived from the neonatal foreskin of two darkly pigmented donors were purchased from Thermo Fisher Scientific (C2025C, Lot #2077650 and #1875821). HEMs were cultured in Medium 254 supplemented with 1% human melanocyte growth supplement (S-016-5, Cascade Biologics/Thermo Fisher Scientific) and 100 units/mL penicillin/streptomycin, under standard conditions (37°C and 5% CO_2_), for up to 15 passages. HEMs were transfected using Nucleofector^®^ Program U-024 for Human Melanocytes (VPD-1003, Basel, Switzerland) according the manufacturer’s protocol.

#### Human pigmented melanoma cells (MNT1)

were a generous gift from Dr. Michael S. Marks (Children’s Hospital of Philadelphia and University of Pennsylvania, Philadelphia, PA) and cultured under standard conditions (37°C and 5% CO_2_) in Dulbecco’s Modified Eagle Medium (DMEM) supplemented with 18% fetal bovine serum (FBS; S11550, Atlanta Biologicals, Flowery Branch, GA), 10% AIM-V™, and 100 units/mL penicillin/streptomycin. For pH experiments, MNT1 cells were transiently transfected with 2 μg plasmid DNA per construct using the Nucleofector^®^ Program U-018 from the Nucleofector Kits for Human Melanocytes and imaged 24 hr following transfection. For melanin assays involving ectopic expression of CLC7 and OCA2, MNT1 cells were pretreated with 300 μM tyrosinase inhibitor propylthiouracil (PTU; P7629, Sigma-Aldrich, St. Louis, MO) for 14 days prior to nucleofection and then released from PTU treatment on the day of transfection. MNT1 cells were collected by cell lysis for melanin assay 5 days following transfection. Transfection efficiency of the CLC7 and OCA2 constructs was approximately 60-70%. For immunostaining, MNT1 cells were transiently transfected 24 hr prior to fixation with 1 μg plasmid DNA per construct using Magnetofection™ (PG60100, OZ Biosciences, San Diego, CA) according to the manufacturer’s recommendations.

#### HeLa cells

(CCL-2, ATCC^®^, Manassas, VA) were cultured under standard conditions (37°C and 5% CO_2_) in DMEM supplemented with 10% FBS and 100 units/ml penicillin/streptomycin. For live-cell experiments involving LysoSensor, HeLa cells were transiently transfected with 2 μg plasmid DNA per construct using the Nucleofector^®^ Program I-013 from the Nucleofector Kits for Human Melanocytes and imaged 24 hr following transfection. For immunostaining, HeLa cells were transiently transfected 24 hr prior to fixation with 1 μg plasmid DNA per construct using Magnetofection™ according to the manufacturer’s recommendations.

#### Lentiviral transduction

shRNAs targeting CLC7, OSTM1, or scrambled (non-targeting, NT) were used to generate lentiviral particles in HEK293FT cells, as previously described (13). Briefly, low passage HEK293FT cells maintained in DMEM with 10% FBS and 100 units/ml penicillin/streptomycin under standard conditions (37°C and 5% CO_2_) were transfected with individual shRNA in pLKO.1 (Broad Institute, Cambridge, MA) mixed with packaging plasmid pCMV-dR8.91 (G193486, GenScript, Piscataway, NJ) and envelope plasmid pVSV-G (G178153, GenScript) at a 10:10:1 ratio using Lipofectamine™ 2000 according to the manufacturer’s recommendations. Viral particles were collected 48 h following transfection and stored at −80°C for 48 hr prior to transduction. Adherent HEMs or MNT1 cells were incubated with viral particles and 8 μg/ml polybrene (H9268, Sigma-Aldrich) for 24 h, then selected for 21 days with 0.6 μg/ml and 1 μg/ml puromycin (Invivo-Gen, San Diego, CA), respectively. The CLC7- and OSTM1-targeting shRNA plasmids were part of the shRNA collection selected for the screen described in (13): CLC7-shRNA #1: 5’-CTCCTCGTCTAGGTTTCTTTA, CLC7-shRNA #2: 5’-AGTATGAGAGCTTGGACTATG, OSTM1-shRNA: 5’-TGCAATGAACATCACTCGAAA.

All cell lines used in this study were routinely tested for mycoplasma contamination using the Hoechst nuclear stain and/or Molecular Probes MycoFluor Mycoplasma Detection Kit (M7006).

### DNA constructs

The plasmid-based expression construct pcDNA4/TO-GFP-CLC7 was generated by amplifying the CLC7 insert of pCMV6-CLC7-Myc-FLAG (RC203450, Origene, Rockville, MD) using PCR with a forward primer (5’-CATCATAAGCTTGGAGCTATGG CCAACGTCTCTAAGAAGGTGTC) containing a Hind III restriction site and Gly-Ala linker at the 5’ end and a reverse primer (5’-CATCATCTCGAGTCACGTCTGGGCCAGCGAG) containing a stop codon and XhoI restriction site at the 3’ end. The CLC7 fragment was then subcloned into the pcDNA/4TO-GFP vector previously generated by inserting eGFP in the AflII/HindIII sites of pcDNA4/TO. For pcDNA4/TO-OSTM1-Myc, the OSTM1-Myc insert from pCMV6-OSTM1-MYC-FLAG (RC209871, Origene) was amplified by PCR using forward primers (OSTM1: 5’-CATCATCTCGAGATGGAGCCGGGCCCGAC) and reverse primers (Myc-STOP: 5’-CATCATTCTAGACTACAGATCCTCTTCTGAGATGAGTTTCTGCTC) containing the XhoI/XbaI restriction sites at the 5’ or 3’ ends, respectively. The OSTM1-Myc fragment was then subcloned into the XhoI and HindIII restriction sites of pcDNA4/TO.

pcDNA4/TO-GFP-CLC7-Y715C mutation was generated by site-directed mutagenesis of the pcDNA4/TO-GFP-CLC7 expression vector using the Q5™ Site-Directed Mutagenesis Kit (New England Biolabs, Ipswich, MA) and complementary primer pairs (forward: 5’-CCGAGACGCCTGCCCGCGCTTCC, reverse: 5’-AAGTCCTTCAGCCTCAGGCGCC). Using the pcDNA4/TO-GFP-CLC7-Y715C expression construct, the CLC7-Y715C insert was PCR amplified and subcloned into the HindIII and XhoI restriction sites of pcDNA4/TO. The N-terminal iRFP tag of piRFP-N3-TYR [plasmid #80152, Addgene, Watertown, MA; (5) was synthesized by GenScript and subsequently subcloned into the respective restriction sites of pcDNA4/TO-CLC7-Y715C. All expression vectors were validated by sequencing (GENEWIZ, South Plainfield, NJ).

The genetically encoded pH indicator RpHiMEL was constructed and validated by GenScript. Briefly, human OCA2 (hOCA2, NM_000275) containing V443I mutations was inserted between the BamHI and XhoI sites of pcDNA4/TO as previously described (6). pH-insensitive iRFP was fused to the N-terminus of the OCA2 V443I mutant using the AfIII and HindIII sites while pH-sensitive Nectarine was inserted in the luminal loop between the first and second transmembrane domains using the BamHI and XbaI sites.

The plasmid-based expression constructs pcDNA4/TO-mCh-OCA2 and pcDNA4/TO-mCh-OCA2-V443I encoding wild type or mutant hOCA2 with an N-terminal mCh tag were generated using the HindIII and XhoI restriction sites as previously described (6).

### Melanin Assays

#### High throughput Melanin Assay

High throughput measurements of melanin content in MNT1 cells expressing CLC7 or OSTM1 shRNA were performed as previously described (13). Briefly, the total number of MNT1 cells seeded in a 96-well plate was determined by taking high contrast brightfield images of each well using the Opera Phenix High-Content Screening System (PerkinElmer, Waltham, MA) and default phase-contrast program. The Opera Phenix analysis software was then used to determine the total number of cells per well for normalizing melanin content. For measuring melanin content, MNT1 cells were lysed with 1N NaOH (S5881, Sigma-Aldrich), left to shake overnight at room temperature, and incubated at 85°C for 30 min to solubilize the melanin. The concentration of melanin per well was determined by spectrophotometric analysis (NanoDrop™ 2000) at an absorbance wavelength of 405 nm and a standard curve of synthetic melanin (M0418, Sigma-Aldrich). The cellular melanin concentration was then calculated as total melanin/cell number. At least three technical replicates and three biological replicates were performed for each sample. Statistical significance was determined by paired two-sample t-test.

#### Classic Melanin Assay

HEMs or MNT1 cells seeded in a 6-well plate at 50-80% confluency were lysed in 1% Triton X-100 (T8787, Sigma-Aldrich) and centrifuged at 14,000 RPM for 30 min at 4°C to separate solubilized protein from melanin. For melanin solubilization, melanin pellets were resuspended in 1N NaOH and incubated at 85 °C for 1 hr. The concentration of melanin per sample was determined by spectrophotometric analysis (NanoDrop™ 2000) at an absorbance wavelength of 405 nm and a standard curve of synthetic melanin. The concentration of protein per sample was determined using the Bicinchoninic Acid (BCA) Protein Assay Kit (23228, Pierce/Thermo Fisher Scientific) according to the manufacturer’s recommendations. The volume of soluble melanin and protein were measured for each sample to calculate total melanin/protein. At least three technical replicates and four biological replicates were performed for each sample. Statistical significance was determined by paired two-sample t-test or one-way ANOVA with Tukey’s multiple comparisons test.

#### Quantification of eumelanin and pheomelanin

HEMs expressing CLC7- or OSTM1-targeting shRNA were seeded in 6-well plates and cultured to 50-80% confluency. Cells were then detached with 0.05% trypsin, suspended in appropriate cell culture media, and lyophilized for subsequent HPLC analysis. Permanganate oxidization, hydroperoxide oxidation, and hydriodic acid hydrolysis were performed on triplicate samples per condition to measure pyrrole 2,3,5-tricarboxylic acid (PTCA), thiazole-2,4,5-tricarboxylic acid (TTCA), and 4-amino-3-hydroxyphenylalanine (4-AHP), respectively (29, 30, 40). The protein concentration of each sample was determined by Lowry-Folin methods to which the PTCA, TTCA and 4-AHP contents were normalized. To correlate eumelanin to PTCA content and pheomelanin to TTCA and 4-AHP content the following conversion factors were applied: eumelanin, PTCA x 38; benzothiazole-type pheomelanin, TTCA × 34, and benzothiazine-type pheomelanin, 4-AHP x 9 (28–30, 40).

### Real-time quantitative polymerase chain reaction (qPCR)

Paired HEMs and MNT1 cells used for classical melanin assays were subjected to qPCR analysis to quantify CLC3, CLC5, CLC7, and OSTM1 mRNA levels. Total RNA was extracted using the RNeasy Plus Mini Kit (74134, Qiagen, Germantown, MD) and reversed transcribed using the iScript™ cDNA Synthesis Kit (1708841, Bio-Rad, Hercules, CA) according to the manufacturer’s instructions. qPCR reactions contained 1 μg cDNA, iTaqTM Universal SYBR Green supermix (1725121, Bio-Rad), and Bio-Rad primers specific to CLC3 (qHsaCED0043281), CLC5 (qHsaCID0007268), CLC7 (qHsaCID0017302), GAPDH (qHsaCED0038674) or IDT primer specific to OSTM1 (Hs.PT.58.38419168) and were performed using the Bio-Rad CFX96 Touch Real-Time PCR Detection System with default cycling parameters. Normalized mRNA expression was calculated using the CFX Maestro Software with GAPDH as the reference gene. Three technical replicates and three biological replicates were performed for each sample. Statistical significance was determined by paired two-sample t-test.

### Western blot

Confluent cells seeded on 35 mm cell culture dishes were rinsed with cold phosphate-buffered saline (PBS) and then lysed using a cold solution of 300mM NaCl, 50mM Tris-HCl (pH 7.4), 1% Triton X-100, and protease inhibitors (5892970001, Sigma-Aldrich). Cell lysates were collected by scrapping and homogenization using a 22-gauge syringe needle and then rotated continuously for 1 h at 4 °C. To remove cell debris and melanin, cell lysates were centrifuged at 14,000 RPM for 30 min at 4 °C. The concentration of protein per sample was determined using the Pierce BCA Protein Assay Kit according to the manufacturer’s recommendations. 25 μg protein from each sample was mixed with LDS Sample Buffer (M00676, GenScript) and loaded in each well of a 4-20% SurePAGE™ Bis-Tris gel (M00656, GenScript). Proteins were separated by SDS-PAGE and transferred to a nitrocellulose membrane (1620145, Bio-Rad). Membranes were blocked with 5% milk in Tris-buffered saline with 0.5% Tween 20 (TBST) for 1 h at room temperature. Immunoblot analysis was performed using mouse monoclonal anti-CLC3 (1:4000; sc-390010, Santa Cruz Biotechnology, Dallas, TX), rabbit polyclonal anti-CLC5 (1:4000; PA5-64891, Thermo Fisher Scientific), rabbit polyclonal anti-CLC7 (1:4000; AP11863b, Abgent, San Diego, CA) or mouse monoclonal anti-GAPDH (1:10,000; AM4300, Thermo Fisher Scientific) and horseradish peroxidase-conjugated goat anti-mouse or anti-rabbit secondary antibodies (1: 10,000 GAPDH, 1:4000 CLC3, 1:8000 CLC5 and CLC7). Immunoblots were visualized using SuperSignal™ West Pico Plus Chemiluminescent Substrate (34579, Thermo Fisher Scientific) and a Bio-Rad Universal Hood II Molecular Imager with CFW-1312M camera. Western blot densitometry band quantification was performed using ImageJ/FIJI (41). Acquired images were converted to 8-bits and percent peak intensity area measurements were obtained for selected bands using the GelAnalyzer plugin. These measurements were used for normalization of CLC3, CLC5, and CLC7 protein expression to GAPDH protein expression (loading control). At least three independent biological replicates were performed for Western blot experiments unless otherwise specified. Statistical significance was determined by paired two-sample t-test.

### Immunofluorescence and confocal microscopy

Cells were seeded on glass coverslips coated with a 0.1% Poly-L-lysine solution (P8920, Sigma-Aldrich) and fixed with 4% paraformaldehyde (P6148, Sigma-Aldrich) in PBS for 20 mins at room temperature. Cells were labelled with primary antibodies (25 min at room temperature) and secondary antibodies (60 min at 37°C) diluted in PBS with 0.2% saponin (47036, Sigma-Aldrich), 0.1% bovine serum albumin (BP1600, Thermo Fisher Scientific), and 0.02% sodium azide (S2002, Sigma-Aldrich) as previously described (42). Coverslips were mounted onto glass slides using Vectashield Antifade Mounting Medium (H-1000, Vector Laboratories, Burlingame, CA) and imaged on an inverted Olympus FV3000 confocal microscope equipped with a 60X objective lens (1.3 NA; UPlan Super Apochromat), a resonant scanner, multi-alkali and cooled GaAsP photomultipliers, and FluoView image acquisition software. Primary antibodies used were rabbit polyclonal anti-CLC7 (1:50; AP11863b, Abgent), mouse monoclonal anti-MYC (1:500; MA1-980, Thermo Fisher Scientific), mouse monoclonal anti-TYRP1 (1:50; 917801, Biolegend, San Diego, CA), and mouse monoclonal anti-LAMP1 (1:10; H4A3, Developmental Studies Hydbridoma Bank, Iowa City, IA). Species- or isotype-specific secondary antibodies from goat or donkey were conjugated to Alexa Fluor 405, Alexa Fluor 488, Alexa Fluor 568, or Alexa Fluor 647 (1:2000; Thermo Fisher Scientific). Colocalization analysis of the acquired images was performed using ImageJ/FIJI (v2.0.0), as previously described (7, 26). Briefly, each multichannel image was converted to 8-bits and background subtracted using a rolling ball radius of 10 pixels. Two regions of interest (ROIs) of equal size were randomly selected for colocalization analysis. All channels were then thresholded (background value = 0, organelle value = 255) to generate binary images and analyzed to obtain the total fluorescence intensity of each channel. Using the multiply channels parameter, composite images were generated of two desired channels. To calculate the percent overlap between these single-channels, the total fluorescence of composite images was divided by the total fluorescence of each single-channel. As positive controls, we quantified the percent overlap of cells expressing GFP-CLC7 and stained with α-CLC7 (**Fig. S3A-B**). Likewise, we quantified cells expressing both RpHiMEL (a variant of OCA2-V443I that retains WT OCA2 localization) and WT OCA2 (**Fig. 4B-C**) to estimate maximum percent overlap.

### pH imaging

#### RpHiMEL

MNT1 cells expressing nontargeting (NT) or CLC7-targeting shRNA were transiently transfected with 2 μg RpHiMEL 24 h prior to pH imaging experiments. Cells seeded on glass coverslips (GG-25, neuVitro, Vancouver, WA) were imaged in extracellular Ringer’s solution (135 mM NaCl, 5 mM KCl, 2 mM CaCl_2_, 2 mM MgCl_2_, 20 mM HEPES, 10 mM D-glucose, pH 7.4) using an Olympus IX71 inverted fluorescence microscope equipped with a 60x objective lens (1.4 NA; UPlanFLN), Hamamatsu EM-CCD digital camera and MetaMorph image acquisition software. Sequential multichannel images were acquired every 10 s for 400 s using excitation and emission wavelengths for Nectarine (λ_ex_ = 540 – 580 nm, λ_em_ = 593 – 668 nm) and for iRFP (λ_ex_ = 588 – 653 nm, λ_em_ = 652 – 717 nm). Baseline fluorescence of cells expressing RpHiMEL was measured for 100 s prior to equilibration. The pH of RpHiMEL-expressing organelles was determined using a calibration curve resulting from equilibration by adding a solution of 24 μM monensin sodium salt (M5273, Sigma-Aldrich), 20 μM tributyltin chloride (T50202, Sigma-Aldrich), 10 μM nigericin sodium salt (N7143, Sigma-Aldrich) in extracellular Ringer’s solution of desired pH [pH 5.0 – 8.0] to cells at the 100 s timepoint. Five independent biological replicates measurements were performed for luminal pH in MNT1 cells expressing NT or CLC7 shRNA. At least three independent biological replicates were performed for calibration at pH 5.0, 6.0, 7.0, and 8.0 in MNT1 cells expressing NT shRNA.

Fluorescence intensity data for RpHiMEL-expressing organelles was acquired in ImageJ/FIJI (v2.0.0). using background-subtracted interleaved image stacks and the Manual Tracking (v2.0) plugin with application of the 2D centring correction (41). Only the RpHiMEL-expressing organelles that remained in the focal plane throughout the time-lapse duration were manually tracked, which resulted in approximately 20 individual organelle measurements per cell. RpHiMEL fluorescence was then calculated as the ratio of the pH-sensitive nectarine fluorescence intensity to the pH-insensitive fluorescence intensity (F_RpHiMEL_ = F_Nectarine_/F_iRFP_) at each timepoint per organelle. To generate the pH calibration curve, the average F_RpHiMEL_ was calculated for each organelle over a 50 s interval after equilibration (300 to 350 s). The calibration curve was fit by linear regression using GraphPad Prism 7 and the average F_RpHiMEL_ was linear across pH 5.0 to 8.0 (R^2^ = 0.99). Using the calibration curve, the pH of individual RpHiMEL-expressing organelles was calculated by averaging the baseline F_RpHiMEL_ from a 30 s interval prior to equilibration (50 to 80 s). Statistical significance was determined by unpaired two-sample t-test.

#### LysoSensor Yellow/Blue DND-160

HeLa and MNT1 cells ectopically expressing mCh-tagged hOCA2 or hOCA2-V443I with or without iRFP-CLC7-Y715C were seeded on glass coverslips (GG-25, neuVitro) for 24 hr prior to live-cell imaging. Cells were incubated for 5 min with 2 μM LysoSensor Yellow/Blue DND-160 (L7545, Invitrogen/Thermo Fisher Scientific) in extracellular Ringer’s solution at 37°C and 5% CO_2_. Following a washing step, cells were imaged in extracellular Ringer’s solution using an Olympus IX71 inverted fluorescence microscope equipped with a 60x objective lens (1.4 NA; UPlanFLN) Hamamatsu EM-CCD digital camera and MetaMorph image acquisition software. For LysoSensor Yellow/Blue DND-160 imaging, single multichannel images were acquired at 60X magnification using wavelength 1 (W1) (λ_ex_ = 352 – 402 nm, λ_em_ = 417 – 477 nm) and wavelength 2 (W2) (λ_ex_ = 354 – 366 nm, λ_em_ = 528 – 556 nm) excitation and emission wavelengths. Paired fluorescence intensity and area data for acidic organelles stained with LysoSensor were manually acquired in ImageJ/FIJI (v2.0.0) using background-subtracted interleaved image stacks of the W1 and W2 channels. CLC7 (WT or Y715C variant) and OSTM1 were co-transfected in all experiments, and only cells expressing all transfected constructs per condition were analyzed. LysoSensor fluorescence was calculated as the ratio of fluorescence intensities (W1/W2). The minimum and maximum thresholds for cross-sectional area measurements were 0.11-25.76 μm^2^ and 0.11-21.93 μm^2^ in HeLa and MNT1 cells, respectively. Live-cell imaging with LysoSensor was performed in at least five individual cells per condition from three independent experiments. Statistical significance was determined by unpaired two-sample t-test or one-way ANOVA with Tukey’s multiple comparisons test. The non-parametric Mann-Whitney test or Kruskal-Wallis test with Dunn’s multiple comparisons test were performed where appropriate.

### Statistical analyses and data visualization

All statistical tests were performed using GraphPad Prism 7. *A priori* statistical analyses were not performed to predetermine sample size. Data distributions and variances were assessed prior to statistical analysis using the parametric paired or unpaired two-sample t-test or one-way ANOVA with Tukey’s multiple comparisons test in GraphPad Prism. Data with unequal variances or non-normal distributions were analyzed using the non-parametric Mann-Whitney test or Kruskal-Wallis test with Dunn’s multiple comparisons test in GraphPad Prism. A p-value of <0.05 was considered statistically significant. All graphs were created using GraphPad Prism 7 and Microsoft Excel (v16.16.25). Figure illustrations were made using Microsoft Powerpoint (v16.16.25) and BioRender (BioRender.com).

## Acknowledgments

We thank members of the Oancea lab for technical support and stimulating discussions. This project was supported by NIH R01AR071382 (to EO), R01AR071382-02S1 (to DCK), and T32GM077995 (to JLS). The authors declare no conflict of interest.

