## Supplemental Figure Legends for "The lysosomal chloride-proton exchanger CLC7 functions in melanosomes as a negative regulator of human pigmentation"

#### Supplementary Figure Legends

**Figure S1. CLC7 downregulation leads to increased pigmentation in MNT1 human melanoma cells.**

**A.** Representative bright field images of MNT1 cells expressing CLC7-targeting shRNA show increased pigmentation compared to cells expressing control non-targeting (NT) shRNA. Scale bar: 20 μm.

**B.** Total cellular melanin concentration (ng melanin/10,000 cells) is increased 2.5-fold and 8.0-fold in MNT1 cells expressing CLC7 shRNA #1 or CLC7 shRNA #2, respectively, compared to NT shRNA cells. ± SEM; n = 2 independent experiments.

**C.** CLC7 mRNA levels were reduced by 78% and 77% in MNT1 cells expressing CLC7 shRNA #1 (dark green) or CLC7 shRNA #2 (light green) compared to control NT shRNA cells (grey). ± SEM; n ≥ 2 independent experiments.

**D.** Compared to MNT1 cells expressing NT shRNA (grey bar), CLC7 protein levels were decreased by 47% and 75% in CLC7 shRNA #1 (dark green) or CLC7 shRNA #2 (light green) cells. *Inset:* Representative immunoblot showing attenuated CLC7 protein levels in MNT1 cells expressing CLC7 shRNA #1 or CLC7 shRNA #2. GAPDH served as a loading control. ± SEM; n = 2 independent experiments.

**E.** Uncropped immunoblot from **Figure 1C** showing decreased CLC7 protein levels in HEMs expressing CLC7 shRNA #1 and CLC7 shRNA #2 compared to NT shRNA cells. GAPDH served as a loading control.

**Figure S2. CLC3 and CLC5 expression in HEMs with CLC7 downregulation.**

**A.** Relative mRNA levels of OSTM1, CLC7, CLC5, and CLC3 in HEMs measured by qPCR. GAPDH served as a reference gene. ± SEM; n = 3 independent experiments.

**B.** Compared to HEMs cells expressing control non-targeting (NT) shRNA, CLC3 mRNA (bar graph) and protein (gel image) levels were not significantly different in CLC7 shRNA expressing cells (P = 0.4725 and P = 0.1601, respectively; Paired two-sample t-test). GAPDH served as a loading control for mRNA (top) and protein (bottom) expression analyses. ± SEM; n = 3 independent experiments.

**C.** Uncropped immunoblot from **B** showing increased CLC3 protein levels in HEMs expressing CLC7 shRNA #1 and CLC7 shRNA #2 compared to NT shRNA cells. GAPDH served as a loading control.

**D.** Compared to HEMs cells expressing NT shRNA, CLC5 protein levels were relatively unchanged in CLC7 shRNA cells (1.4-fold change; P = 0.0880, Paired two-sample t-test). GAPDH served as a loading control. ± SEM; n = 3 independent experiments.

**E.** Uncropped immunoblot from **D** showing increased CLC5 protein levels in HEMs expressing CLC7 shRNA #1 and CLC7 shRNA #2 compared to NT shRNA cells. GAPDH served as a loading control.

**Figure S3. GFP-CLC7 colocalizes with endogenous CLC7 and mCherry-OCA2.**

**A.** HEMs transiently transfected with GFP-CLC7 and OSTM1-Myc. Endogenous CLC7 and OSTM1-Myc were detected by anti-CLC7 (α-CLC7) or anti-Myc (α-Myc) immunostaining, respectively. *Inset:* Magnified view of boxed areas (white). Arrows indicate organelles coexpressing GFP-CLC7, OSTM1-Myc, and endogenous CLC7. Scale bar: 10 µm.

**B.** Quantification of **A** showing reciprocal overlap between endogenous CLC7 and GFP-CLC7 in HEMs. Endogenous CLC7 colocalizes with 58% of total GFP-CLC7. GFP-CLC7 colocalizes with 85% of total endogenous CLC7. ± SEM; n = 4 independent experiments.

**C.** HeLa cells transiently transfected with GFP-CLC7, OSTM1-Myc, and mCh-OCA2. OSTM1-Myc was observed by anti-Myc (α-Myc) immunostaining. *Inset:* Magnified view of boxed areas (white). Arrows indicate organelles coexpressing CLC7, OSTM1, and OCA2. Scale bar: 10 µm.

**D.** Quantification of **C** showing reciprocal overlap between overexpressed GFP-CLC7 and mCh-OCA2 (top) or GFP-CLC7 and OSTM1-Myc (bottom) in HeLa cells. mCh-OCA2 and OSTM1-Myc colocalize with ≥72% of total GFP-CLC7. GFP-CLC7 colocalizes with ≥77% of total mCh-OCA2 or OSTM1-Myc. ± SEM; n = 6 independent experiments.

**Figure S4. OCA2 expression in MNT1 cells expressing CLC7-Y715C rescues vacuole size and acidic pH.**

**A.** MNT1 cells transiently transfected with iRFP-CLC7-Y715C and OSTM1-Myc together with mCh-OCA2-V443I (top) or mCh-OCA2-WT (bottom). OSTM1-Myc was observed by anti-Myc (α-Myc) immunostaining. Inset: Magnified view of boxed areas (white). Arrows indicate organelles coexpressing CLC7, OSTM1, and OCA2. Scale bar: 10 µm.

**B.** Average cellular organelle area (in $\mu m^2$) obtained as the average of LysoSensor-stained organelles from individual MNT1 cells expressing the indicated proteins. The average organelle area per MNT1 cell overexpressing mCh-OCA2-WT or mCh-OCA2-V443I is 0.78 ± 0.06 $\mu m^2$ and 0.80 ± 0.07 $\mu m^2$, respectively (± SEM; P = 0.8763, Mann-Whitney test). Mean ± SEM.

**C.** Area values of individual LysoSensor-stained organelles from MNT1 cells overexpressing mCh-OCA2-V443I (grey circles) or mCh-OCA2-V443I (green circles). Compared to control cells expressing OCA2-V443I, the average organelle area is unchanged in cells expressing OCA2-WT (P = 0.5019, Mann-Whitney test). Mean ± SEM.

**D.** Average cellular $I_{W1}/I_{W2}$ obtained as the average of LysoSensor-stained organelles from individual MNT1 cells expressing the indicated proteins. The average organelle $I_{W1}/I_{W2}$ per MNT1 cell overexpressing mCh-OCA2-WT or mCh-OCA2-V443I is 0.14 ± 0.02 and 0.18 ± 0.03, respectively (± SEM; P = 0.3464, Unpaired two-sample t-test). Box plots show the data medians, $25^{th}$ and $75^{th}$ percentiles, and extrema.

**E.** $I_{W1}/I_{W2}$ values of individual LysoSensor-stained organelles from MNT1 cells overexpressing mCh-OCA2-V443I (grey circles) or mCh-OCA2-V443I (green circles). Compared to control cells expressing OCA2-V443I, the average organelle area is decreased 1.27-fold in cells expressing OCA2-WT (P = 0.0002, Mann-Whitney test). Mean ± SEM. Organelle $I_{W1}/I_{W2}$ measurements in **D & E** were paired with organelle area measurements in **B & C**. Area and $I_{W1}/I_{W2}$ measurements were acquired from ≥ 250 organelles per cell from at least 5 cells; n = 3 independent experiments.

**F.** Representative widefield fluorescence images of LysoSensor-stained organelles in MNT1 cells transiently expressing iRFP-CLC7-Y715C and OSTM1-Myc with mCh (empty vector; top) or mCh-OCA2-WT (bottom). LysoSensor dual fluorescence emission $I_{W1}$ (yellow, Lyso-W1) and $I_{W2}$ (blue, Lyso-W2), and the ratio $I_{W1}/I_{W2}$ are shown in pseudocolor, with higher $I_{W1}/I_{W2}$ indicating increased organelle acidity as shown by the pH scale. Scale bar: 10 µm.

**G.** Average cellular organelle area (in $\mu m^2$) obtained as the average of LysoSensor-stained organelles from individual MNT1 cells expressing the indicated proteins. The average organelle area per MNT1 cell overexpressing iRFP-CLC7-Y715C and OSTM1-Myc with mCh-EV or mCh-OCA2-WT is 3.24 ± 0.77 $\mu m^2$ and 0.86 ± 0.11 $\mu m^2$, respectively (± SEM; P = 0.0010, Mann-Whitney test). Mean ± SEM.

**H.** Area values of individual LysoSensor-stained organelles from MNT1 cells overexpressing iRFP-CLC7-Y715C and OSTM1-Myc with mCh-EV (grey circles) or mCh-OCA2-WT (green circles). Compared to control cells with mCh-EV coexpression, the average organelle area is decreased 1.17-fold in cells expressing OCA2-WT (P < 0.0001, Mann-Whitney test). Mean ± SEM.

**I.** Average cellular $I_{W1}/I_{W2}$ obtained as the average of LysoSensor-stained organelles from individual MNT1 cells expressing the indicated proteins. The average organelle $I_{W1}/I_{W2}$ per MNT1 cell overexpressing iRFP-CLC7-Y715C and OSTM1-Myc with mCh-EV or mCherry-OCA2-WT is 0.96 ± 0.02 and 0.61 ± 0.03, respectively (± SEM; P = 0.0667, Unpaired two-sample t-test). Box plots show the data medians, 25$^{th}$ and 75$^{th}$ percentiles, and extrema.

**J.** $I_{W1}/I_{W2}$ values of individual LysoSensor-stained organelles from MNT1 cells overexpressing iRFP-CLC7-Y715C and OSTM1-Myc with mCh-OCA2-V443I (grey circles) or mCh-OCA2-V443I (green circles). Compared to control cells expressing OCA2-V443I, the average organelle area is decreased 1.61-fold in cells expressing OCA2-WT (P < 0.0001, Mann-Whitney test). Mean ± SEM. Organelle $I_{W1}/I_{W2}$ measurements in **I & J** were paired with organelle area measurements in **G & H**. Area and $I_{W1}/I_{W2}$ measurements were acquired from ≥ 371 organelles per cell from at least 5 cells; n = 3 independent experiments.

**Figure S5. Expression of CLC7-Y715C results in enlarged and acidified cytosolic vacuoles in HeLa cells.**

**A.** HeLa cells transiently transfected with iRFP-CLC7-Y715C and OSTM1-Myc show enlarged cytosolic vacuoles. Vacuoles were observed by bright field (BF) microscopy. OSTM1-Myc was observed by anti-Myc (α-Myc) immunostaining. Scale bar: 10 µm.

**B.** LysoSensor accumulates in cytosolic vacuoles observed in mCh-expressing HeLa cells transiently transfected with iRFP-CLC7-Y715C and OSTM1-Myc. HeLa cells were incubated with LysoSensor Yellow/Blue DND-160 and imaged using widefield fluorescence microscopy. Expression of mCherry in HeLa cells was used to facilitate the visualization of cytosolic vacuoles. Cytosolic vacuoles observed in the mCh-expressing cells were stained with LysoSensor and analyzed using the yellow fluorescent emission $I_{W1}$ (Lyso-W1). Scale bar: 10 µm.

**C.** Representative widefield fluorescence images of LysoSensor-stained organelles in HeLa cells transiently expressing OSTM1-Myc with iRFP-CLC7-Y715C (top) or iRFP-CLC7-WT (wild type, bottom). LysoSensor dual fluorescence emission $I_{W1}$ (yellow, Lyso-W1) and $I_{W2}$ (blue, Lyso-W2), and the ratio $I_{W1}/I_{W2}$ are shown in pseudocolor, with higher $I_{W1}/I_{W2}$ indicating increased acidity as shown by the pH scale. Scale bar: 10 μm.

**D.** Average cellular organelle area (in $\mu m^2$) obtained as the average of LysoSensor-stained organelles from individual HeLa cells expressing the indicated proteins. The average organelle area per HeLa cell transiently expressing OSTM1-Myc with iRFP-CLC7-WT or iRFP-CLC7-Y715C is 0.63 ± 0.02 $\mu m^2$ and 5.49 ± 1.36 $\mu m^2$, respectively (± SEM; P = 0.0010, Mann-Whitney test). Mean ± SEM.

**E.** Area values of individual LysoSensor-stained organelles from HeLa cells overexpressing OSTM1-Myc with iRFP-CLC7-WT (grey circles) or iRFP-CLC7-Y715C (green circles). The average organelle area in cells with CLC7-Y715C is increased 7.8-fold compared to CLC7-WT cells (P < 0.0001, Mann-Whitney test), indicating that CLC7-Y715C overexpression results in increased organelle size. Mean ± SEM.

**F.** Frequency distribution of organelle area from values obtained in **E**. CLC7-Y715C expression shifts the organelle area distribution to higher values, indicating increased organelle size in HeLa cells expressing CLC7-Y715C compared to CLC7-WT (P < 0.0001, Kolmogorov-Smirnov test).

**G.** Average cellular $I_{W1}/I_{W2}$ obtained as the average of LysoSensor-stained organelles from individual HeLa cells expressing the indicated proteins. The average organelle $I_{W1}/I_{W2}$ per HeLa cell transiently expressing OSTM1-Myc iRFP-CLC7-WT or iRFP-CLC7-Y715C is 0.21 ± 0.02 and 1.51 ± 0.19, respectively (± SEM; P = 0.0010, Mann-Whitney test). A higher $I_{W1}/I_{W2}$ indicates increased organelle acidity as shown by the pH scale (right). Box plots show the data medians, 25$^{th}$ and 75$^{th}$ percentiles, and extrema.

**H.** $I_{W1}/I_{W2}$ values of individual LysoSensor-containing organelles from HeLa cells overexpressing OSTM1-Myc with iRFP-CLC7-WT (grey circles) or iRFP-CLC7-Y715C (green circles). The average organelle $I_{W1}/I_{W2}$ in cells with CLC7-Y715C is increased 6.9-fold compared to CLC7-WT cells, indicating that CLC7-Y715C increases organelle acidity (P < 0.0001, Mann-Whitney test). Mean ± SEM.

**I.** Frequency distribution of organelle $I_{W1}/I_{W2}$ from values obtained in **H**. CLC7-Y715C expression shifts the organelle $I_{W1}/I_{W2}$ distribution to higher values, indicating increased organelle acidity in HeLa cells expressing CLC7-Y715C compared to CLC7-WT (P < 0.0001, Kolmogorov-Smirnov test). Organelle $I_{W1}/I_{W2}$ measurements in **G-I** were paired with organelle area measurements in **D-F**, obtained from at least 113 organelles from ≥ 5 cells were analyzed; n = 5 independent experiments.

**Figure S6. OCA2 rescues cytosolic vacuole size and pH in HeLa cells expressing CLC7-Y715C.**

**A.** HeLa cells transiently transfected with iRFP-CLC7-Y715C and OSTM1-Myc together with mCh-OCA2-V443I (top) or mCh-OCA2-WT (bottom). Bright field (BF) was used to visualize cytosolic vacuoles in transfected cells. OSTM1-Myc was observed by anti-Myc (α-Myc) immunostaining. Inset: Magnified view of boxed areas (white). Arrows indicate organelles coexpressing CLC7, OSTM1, and OCA2. Scale bar: 10 μm.

**B.** Average cellular organelle area (in $\mu m^2$) obtained as the average of LysoSensor-stained organelles from individual HeLa cells expressing the indicated proteins. The average organelle area per HeLa cell overexpressing mCh-OCA2-WT or mCh-OCA2-V443I is 0.82 ± 0.07 $\mu m^2$ and 0.77 ± 0.07 $\mu m^2$, respectively (± SEM; P = 0.6056, Unpaired two-sample t-test). Mean ± SEM.

**C.** Area values of individual LysoSensor-stained organelles from HeLa cells overexpressing mCh-OCA2-V443I (grey circles) or mCh-OCA2-V443I (green circles). Compared to control cells expressing OCA2-V443I, the average organelle area is unchanged in cells expressing OCA2-WT ($P = 0.1483$, Unpaired two-sample t-test). Mean ± SEM.

**D.** Average cellular $I_{W1}/I_{W2}$ obtained as the average of LysoSensor-stained organelles from individual HeLa cells expressing the indicated proteins. The average organelle $I_{W1}/I_{W2}$ per HeLa cell overexpressing mCh-OCA2-WT or mCh-OCA2-V443I is 0.21 ± 0.04 and 0.38 ± 0.05, respectively (± SEM; $P = 0.0169$, Unpaired two-sample t-test). Box plots show the data medians, 25$^{th}$ and 75$^{th}$ percentiles, and extrema.

**E.** $I_{W1}/I_{W2}$ values of individual LysoSensor-stained organelles from HeLa cells overexpressing mCh-OCA2-V443I (grey circles) or mCh-OCA2-V443I (green circles). Compared to control cells expressing OCA2-V443I, the average organelle area is decreased 1.74-fold in cells expressing OCA2-WT ($P < 0.0001$, Mann-Whitney test). Mean ± SEM. Organelle $I_{W1}/I_{W2}$ measurements in **D & E** were paired with organelle area measurements in **B & C**. Area and $I_{W1}/I_{W2}$ measurements were acquired from ≥ 464 organelles per cell from at least 11 cells; n = 3 independent experiments.

**F.** Representative widefield fluorescence images of LysoSensor-stained organelles in HeLa cells transiently expressing iRFP-CLC7-Y715C and OSTM1-Myc with mCh-EV (empty vector; top) or mCh-OCA2-WT (bottom). LysoSensor dual fluorescence emission $I_{W1}$ (yellow, Lyso-W1) and $I_{W2}$ (blue, Lyso-W2), and the ratio $I_{W1}/I_{W2}$ are shown in pseudocolor, with higher $I_{W1}/I_{W2}$ indicating increased organelle acidity as shown by the pH scale. Scale bar: 10 µm.

**G.** Average cellular organelle area (in µm$^2$) obtained as the average of LysoSensor-stained organelles from individual HeLa cells expressing the indicated proteins. The average organelle area per HeLa cell overexpressing iRFP-CLC7-Y715C and OSTM1-Myc with mCh-EV or mCh-OCA2-WT is 7.71 ± 1.20 µm$^2$ and 0.97 ± 0.38 µm$^2$, respectively (± SEM; $P = 0.0040$, Mann-Whitney test). Mean ± SEM.

**H.** Area values of individual LysoSensor-stained organelles from HeLa cells overexpressing iRFP-CLC7-Y715C and OSTM1-Myc with mCh-EV (grey circles) or mCh-OCA2-WT (green circles). Compared to control cells with mCh-EV coexpression, the average organelle area is decreased 9.21-fold in cells expressing OCA2-WT ($P < 0.0001$, Mann-Whitney test). Mean ± SEM.

**I.** Average cellular $I_{W1}/I_{W2}$ obtained as the average of LysoSensor-stained organelles from individual HeLa cells expressing the indicated proteins. The average organelle $I_{W1}/I_{W2}$ per HeLa cell expressing iRFP-CLC7-Y715C and OSTM1-Myc with mCh-EV or mCh-OCA2-WT is 1.69 ± 0.06 and 0.49 ± 0.02, respectively (± SEM; $P = 0.0040$, Mann-Whitney test). Box plots show data medians, 25$^{th}$ and 75$^{th}$ percentiles, and extrema.

**J.** $I_{W1}/I_{W2}$ values of individual LysoSensor-stained organelles from HeLa cells overexpressing iRFP-CLC7-Y715C and OSTM1-Myc with mCh-OCA2-V443I (grey circles) or mCh-OCA2-V443I (green circles). Compared to control cells expressing OCA2-V443I, the average organelle area is decreased 4.23-fold in cells expressing OCA2-WT ($P < 0.0001$, Mann-Whitney test). Mean ± SEM. Organelle $I_{W1}/I_{W2}$ measurements in **I & J** were paired with organelle area measurements in **G & H**. Area and $I_{W1}/I_{W2}$ measurements were acquired from ≥ 134 organelles per cell from at least 4 cells.

### Figure S1.

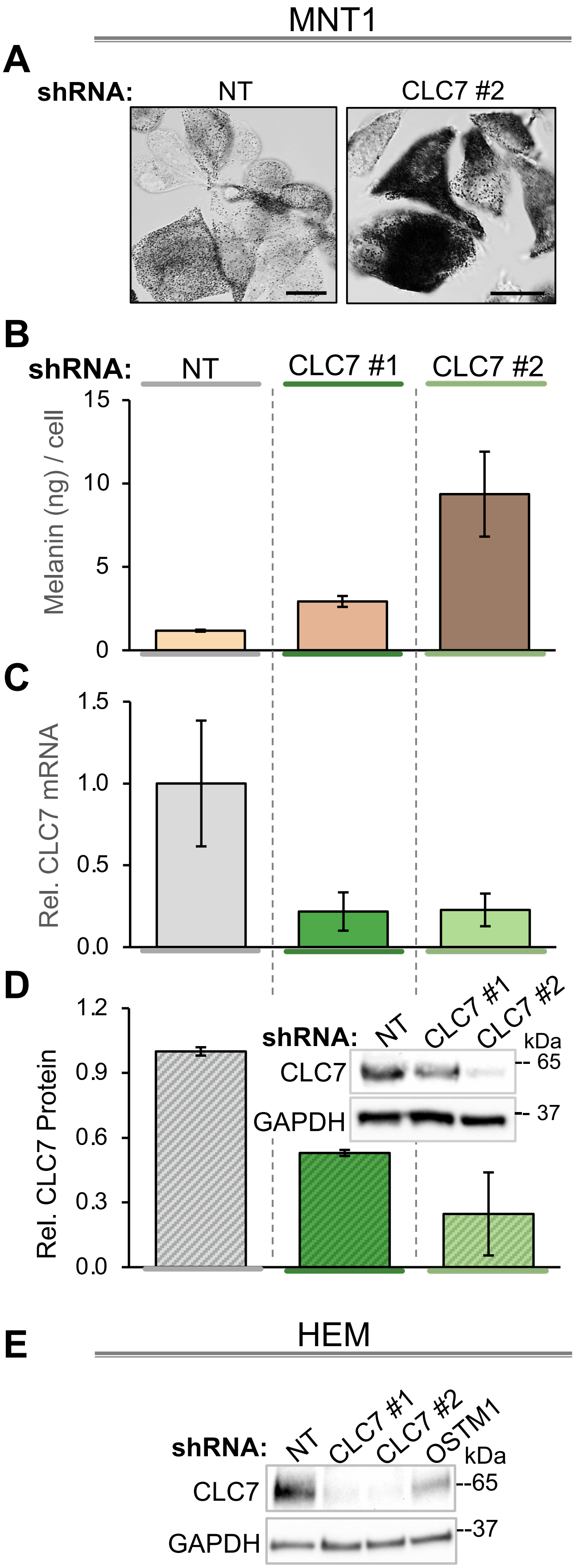
MNT1
A
shRNA:
NT
CLC7 #2
B
shRNA:
NT
CLC7 #1
CLC7 #2
Melanin (ng) / cell
15
10
5
0
C
Rel. CLC7 mRNA
1.5
1.0
0.5
0.0
D
Rel. CLC7 Protein
1.2
0.9
0.6
0.3
0.0
shRNA:
NT
CLC7 #1
CLC7 #2
kDa
CLC7
GAPDH
65
37
E
HEM
shRNA:
NT
CLC7 #1
CLC7 #2
OSTM1
kDa
CLC7
GAPDH
65
37

#### Figure S2.

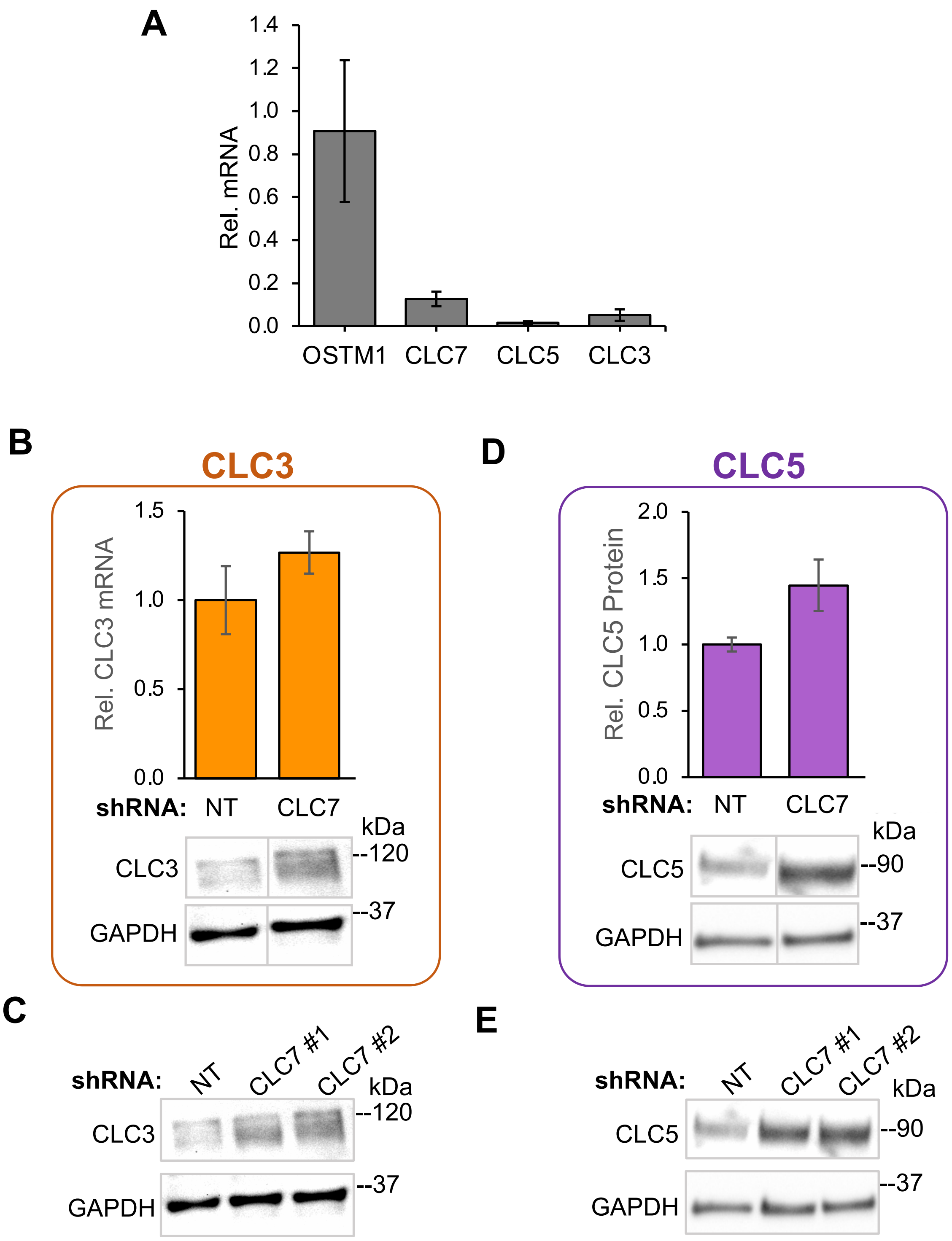
A
1.4
1.2
1.0
0.8
0.6
0.4
0.2
0.0
Rel. mRNA
OSTM1
CLC7
CLC5
CLC3
B
CLC3
1.5
1.0
0.5
0.0
Rel. CLC3 mRNA
shRNA: NT CLC7
kDa
CLC3
--120
--37
GAPDH
D
CLC5
2.0
1.5
1.0
0.5
0.0
Rel. CLC5 Protein
shRNA: NT CLC7
kDa
CLC5
--90
--37
GAPDH
C
shRNA: NT CLC7 #1 CLC7 #2
kDa
CLC3
--120
--37
GAPDH
E
shRNA: NT CLC7 #1 CLC7 #2
kDa
CLC5
--90
--37
GAPDH

**Figure S3.**

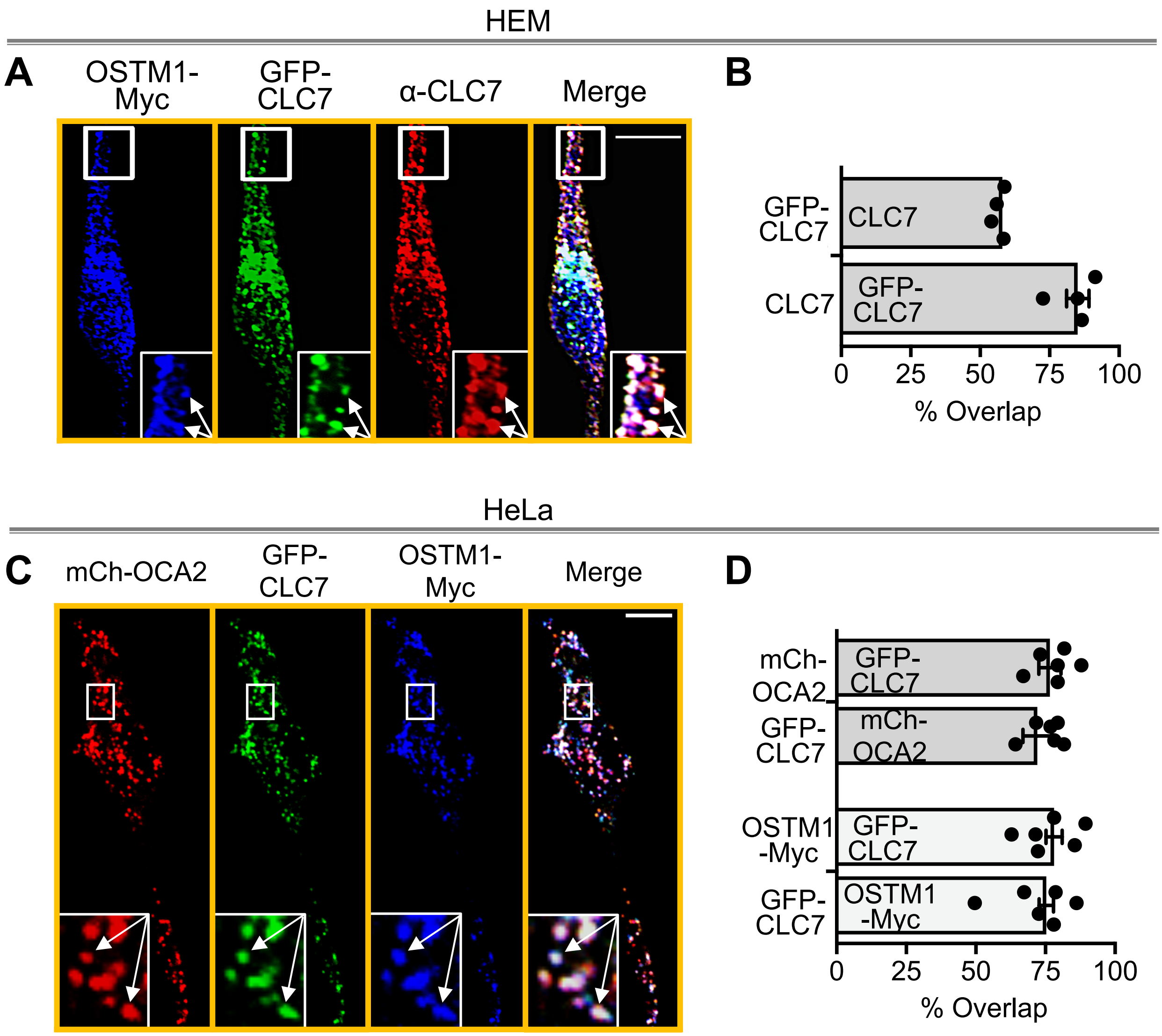
HEM
A
OSTM1-Myc
GFP-CLC7
α-CLC7
Merge
B
GFP-CLC7
CLC7
CLC7
GFP-CLC7
0
25
50
75
100
% Overlap
HeLa
C
mCh-OCA2
GFP-CLC7
OSTM1-Myc
Merge
D
mCh-OCA2
GFP-CLC7
GFP-CLC7
mCh-OCA2
OSTM1-Myc
GFP-CLC7
GFP-CLC7
OSTM1-Myc
0
25
50
75
100
% Overlap

### Figure S4.

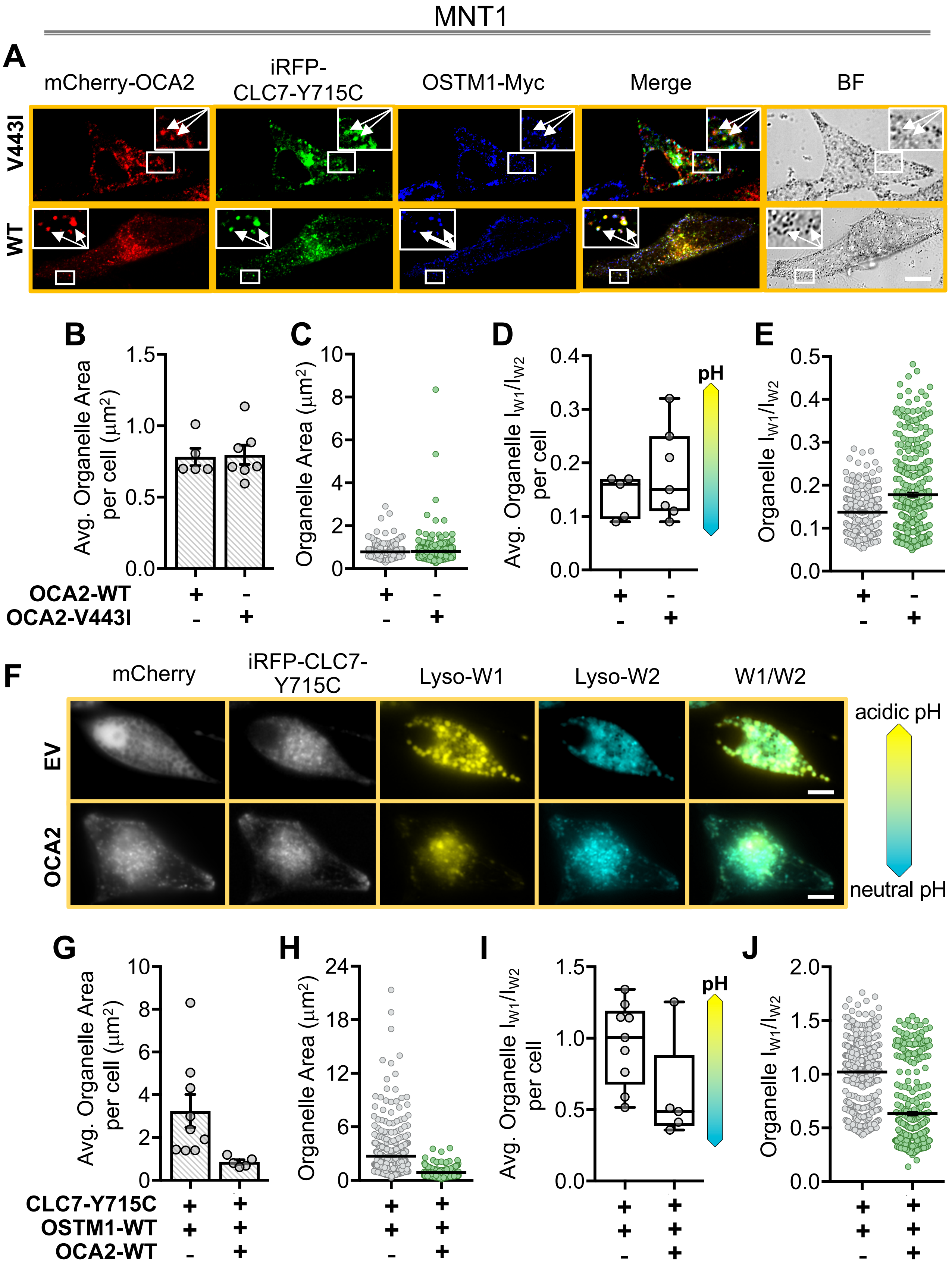

MNT1
A
mCherry-OCA2
iRFP-CLC7-Y715C
OSTM1-Myc
Merge
BF
V443I
WT
B
C
D
E
Avg. Organelle Area per cell (μm²)
Organelle Area (μm²)
Avg. Organelle $I_{W1}/I_{W2}$ per cell
Organelle $I_{W1}/I_{W2}$
pH
OCA2-WT
OCA2-V443I
F
mCherry
iRFP-CLC7-Y715C
Lyso-W1
Lyso-W2
W1/W2
EV
OCA2
acidic pH
neutral pH
G
H
I
J
CLC7-Y715C
OSTM1-WT
OCA2-WT

Figure S5.

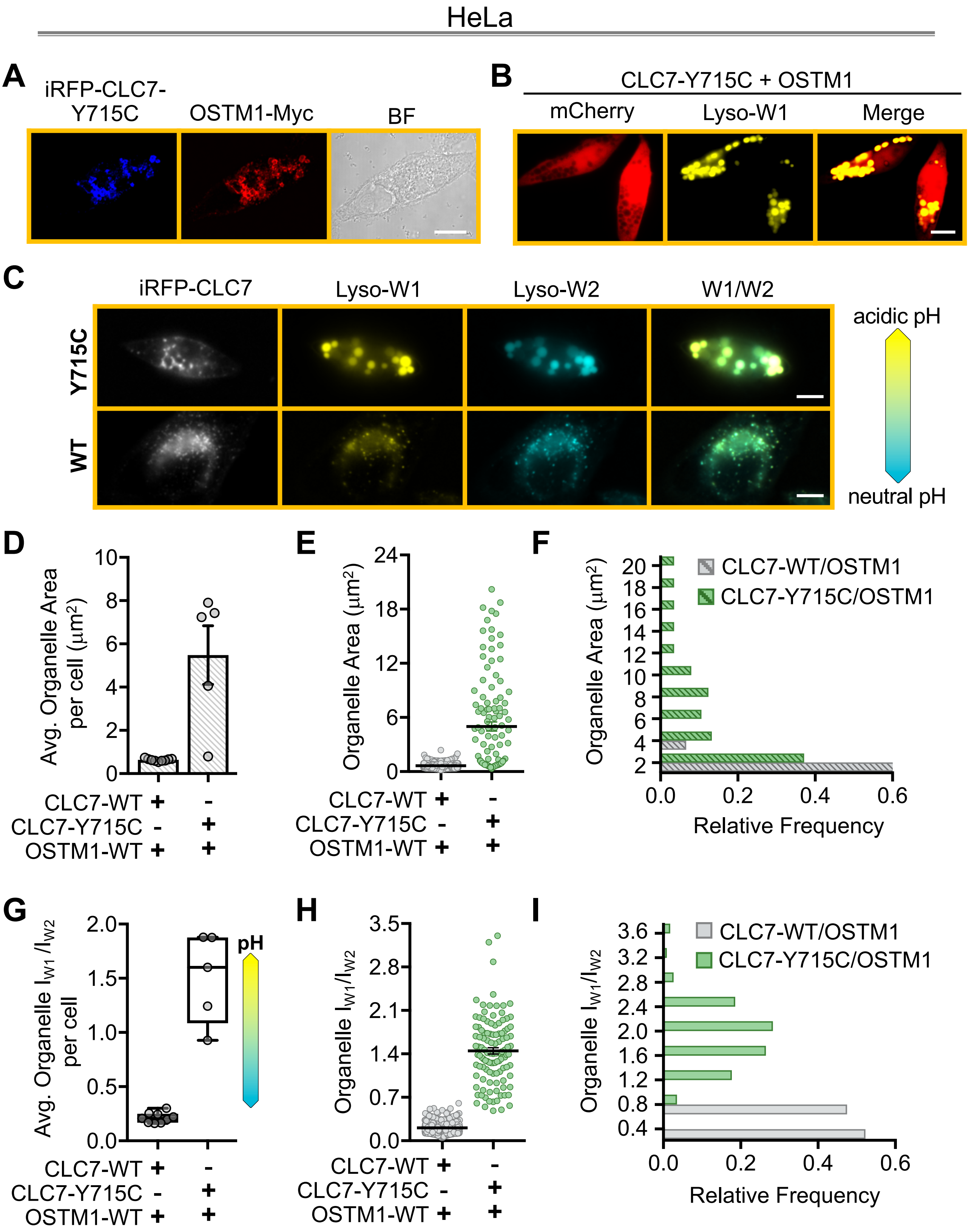
HeLa
A
iRFP-CLC7-Y715C
OSTM1-Myc
BF
B
CLC7-Y715C + OSTM1
mCherry
Lyso-W1
Merge
C
iRFP-CLC7
Lyso-W1
Lyso-W2
W1/W2
Y715C
WT
acidic pH
neutral pH
D
Avg. Organelle Area per cell (μm²)
CLC7-WT + -
CLC7-Y715C - +
OSTM1-WT + +
E
Organelle Area (μm²)
F
Organelle Area (μm²)
CLC7-WT/OSTM1
CLC7-Y715C/OSTM1
Relative Frequency
G
Avg. Organelle I_W1/I_W2 per cell
pH
H
Organelle I_W1/I_W2
I
Organelle I_W1/I_W2
CLC7-WT/OSTM1
CLC7-Y715C/OSTM1
Relative Frequency

### Figure S6.

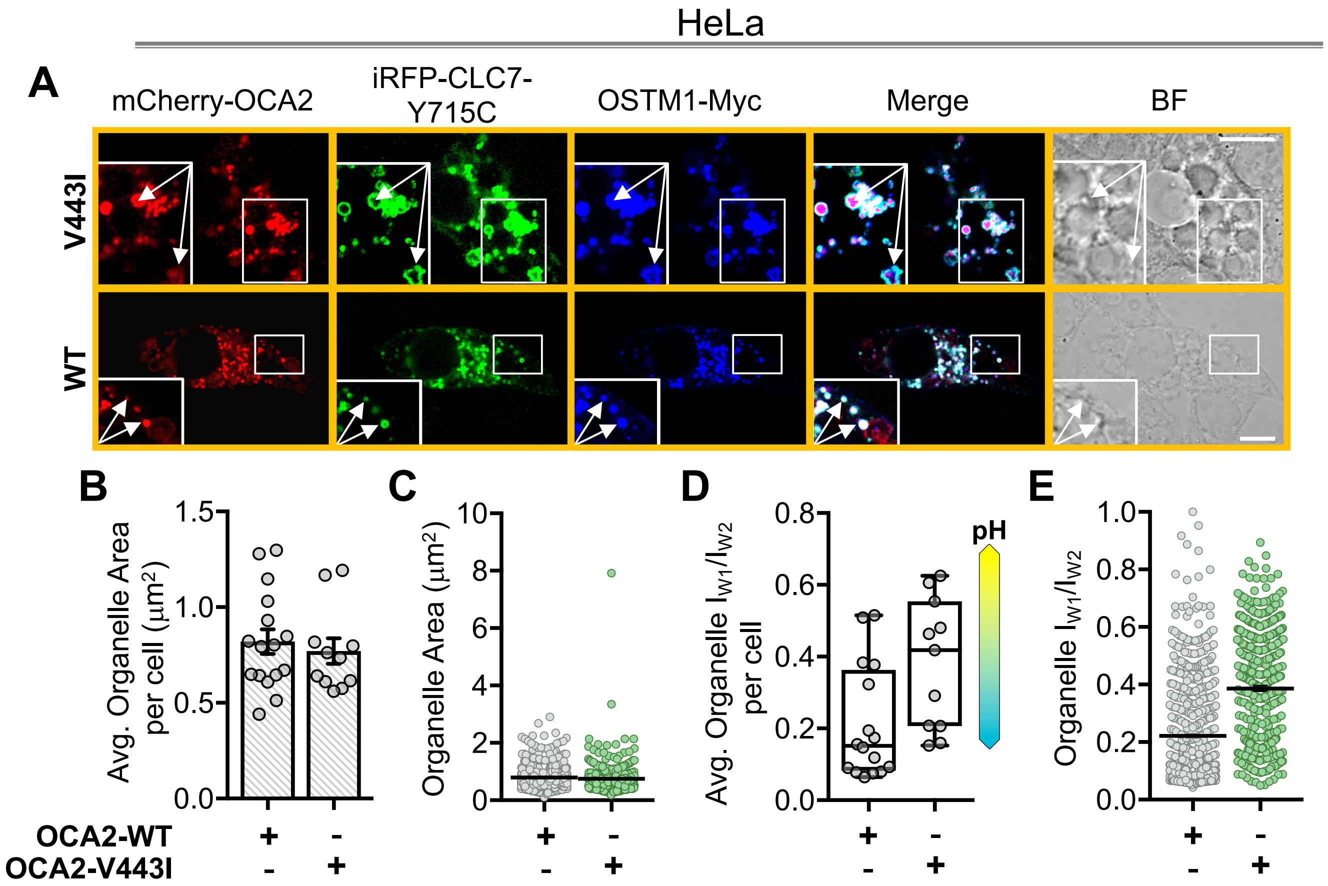
HeLa
A
mCherry-OCA2
iRFP-CLC7-Y715C
OSTM1-Myc
Merge
BF
V443I
WT
B
Avg. Organelle Area per cell (µm2)
1.5
1.0
0.5
0.0
OCA2-WT
OCA2-V443I
+
-
-
+
C
Organelle Area (µm2)
10
8
6
4
2
0
D
Avg. Organelle IW1/IW2 per cell
0.8
0.6
0.4
0.2
0.0
pH
E
Organelle IW1/IW2
1.0
0.8
0.6
0.4
0.2
0.0
